# Obligate cross-feeding of metabolites is common in soil microbial communities

**DOI:** 10.1101/2025.01.29.635426

**Authors:** Ghada Yousif, Francisco Zorrilla, Swagatika Dash, Leonardo Oña, Aditi Shekhar, Samir Giri, Rui Guan, Sharvari Harshe, Michael Itermann, Daphne Welter, Vladimir Benes, Kiran R. Patil, Christian Kost

## Abstract

Many microorganisms are refractory to laboratory cultivation. One possible explanation, known as the great plate count anomaly, is metabolic dependencies among community members. However, systematic studies of these interactions in communities like soil are missing, hindering advances in understanding these ecologically important ecosystems. Here, we address this issue by systematically analysing 6,931 bacterial isolates of 27 soil microbial communities. We find that the growth of up to 50% of all community members depended essentially on supplementation with amino acids, vitamins, or nucleotides. In 73% of cases, supplementation with multiple amino acids was necessary. Genomic analysis of 62 strains revealed that accumulation of insertion sequences and specific gene loss was associated with the observed auxotrophies. Finally, genome-scale metabolic models, computational analyses, and cocultivation experiments demonstrated that other co-occurring genotypes complemented the metabolic needs of auxotrophs, thus facilitating their growth. Our results demonstrate that soil bacteria exist within integrated metabolic networks, which hampers their cultivation.

## Introduction

The great plate count anomaly is a major conundrum in microbiology^1^. This phenomenon describes the observation that the majority of bacterial species, which can be observed under the microscope or that are detectable by sequencing-based approaches, cannot be, or only with great difficulty, cultivated under laboratory conditions^1^. While several hypotheses have been proposed to explain this pattern^2^, metabolic dependencies stand out^3^. The basic idea behind this hypothesis is that the ecological conditions in natural environments favour the evolution of metabolic dependencies among community members^4^. The resulting obligate metabolic relationships may allow strains to survive in their natural habitat, yet thwart their cultivation as monocultures under laboratory conditions when interactions with other community members are disrupted^3,5^.

However, why should bacteria give up their physiological autonomy and become dependent on the provisioning of metabolites by other co-occurring strains? The so-called Black Queen Hypothesis (BQH) provides a mechanistic explanation^4^. The premise of the BQH is that growing bacteria unavoidably release various metabolic byproducts into the extracellular environment^6^. As a consequence, bacterial strains that lose the ability to autonomously produce certain compounds (e.g., amino acids, vitamins, or nucleotides) can take advantage of metabolites that are released by other strains^5^. In addition, ample evidence from numerous laboratory studies demonstrates that mutants, which cannot biosynthesize these compounds, gain fitness advantages when taking up the required metabolite from the environment^5^. In combination, these two observations provide a plausible explanation for the evolution of unidirectional cross-feeding interactions between a prototrophic (i.e., genotype able to produce a given compound) metabolite donor and an auxotrophic (i.e., genotype unable to produce a given compound) recipient. Once emerged, these interactions may become more intertwined as genotypes coevolve, thus facilitating the origin of multipartite interaction networks, which can involve various combinations of auxotrophic and prototrophic genotypes^3^.

Although numerous laboratory-based studies^7–9^ support the BQH, the prevalence of synergistic metabolic interactions in natural microbial communities is frequently questioned^10^. One argument against this idea is mainly based on a theoretical model that suggests the frequent mixing of metabolically dependent genotypes generally selects against obligate cross-feeding^11^. In addition, coculture experiments with pairs of bacterial strains isolated from natural habitats find synergistic interactions between bacterial strains to be rare^12^. However, none of these previous studies explicitly considered the potential significance of interactions that involved auxotrophic genotypes. In addition, knowledge of the prevalence of auxotrophic strains in natural bacterial communities is generally lacking. Thus, a formal verification of the BQH is a topical question in microbial ecology. Such a test should, in particular, (i) quantify the abundance of auxotrophic strains in natural microbial communities, and (ii) verify whether cross-feeding of metabolites between bacterial isolates can explain their existence.

To systematically address this question, we analysed 27 soil bacterial communities to experimentally quantify how many isolates display auxotrophic phenotypes. Moreover, the genomes of a representative set of isolated strains were sequenced to determine the genetic basis of the observed auxotrophic phenotypes. Finally, we combined pairwise and consortium-level coculture experiments with the modelling of metabolic interactions among assembled genomes to validate whether cross-feeding of essential metabolites can explain the occurrence of auxotrophic genotypes in soil microbial communities. As such, our work represents the first test of the BQH in communities of soil-dwelling bacteria.

## Results

### Metabolic auxotrophies are common among culturable soil bacteria

To unravel how prevalent auxotrophic bacterial strains are in soil microbial communities, nine different sites were investigated (Supplementary Fig. 1). From each site, a column of soil was collected to extract three soil particles (Fig. 1a, Methods). Bacteria were isolated on a minimal medium (MMAB) that contained fructose (5 g l^-1^) as the sole carbon source^13^ and to which the following supplementations were added: amino acids, vitamins, and nucleosides (Methods). While we managed to obtain single-cell colonies of many strains, some remained mixed (i.e., 35.7% of 1,421 isolates) and could not be grown individually. Subsequently, these mixed clones were treated as a unit.

**Fig 1.**
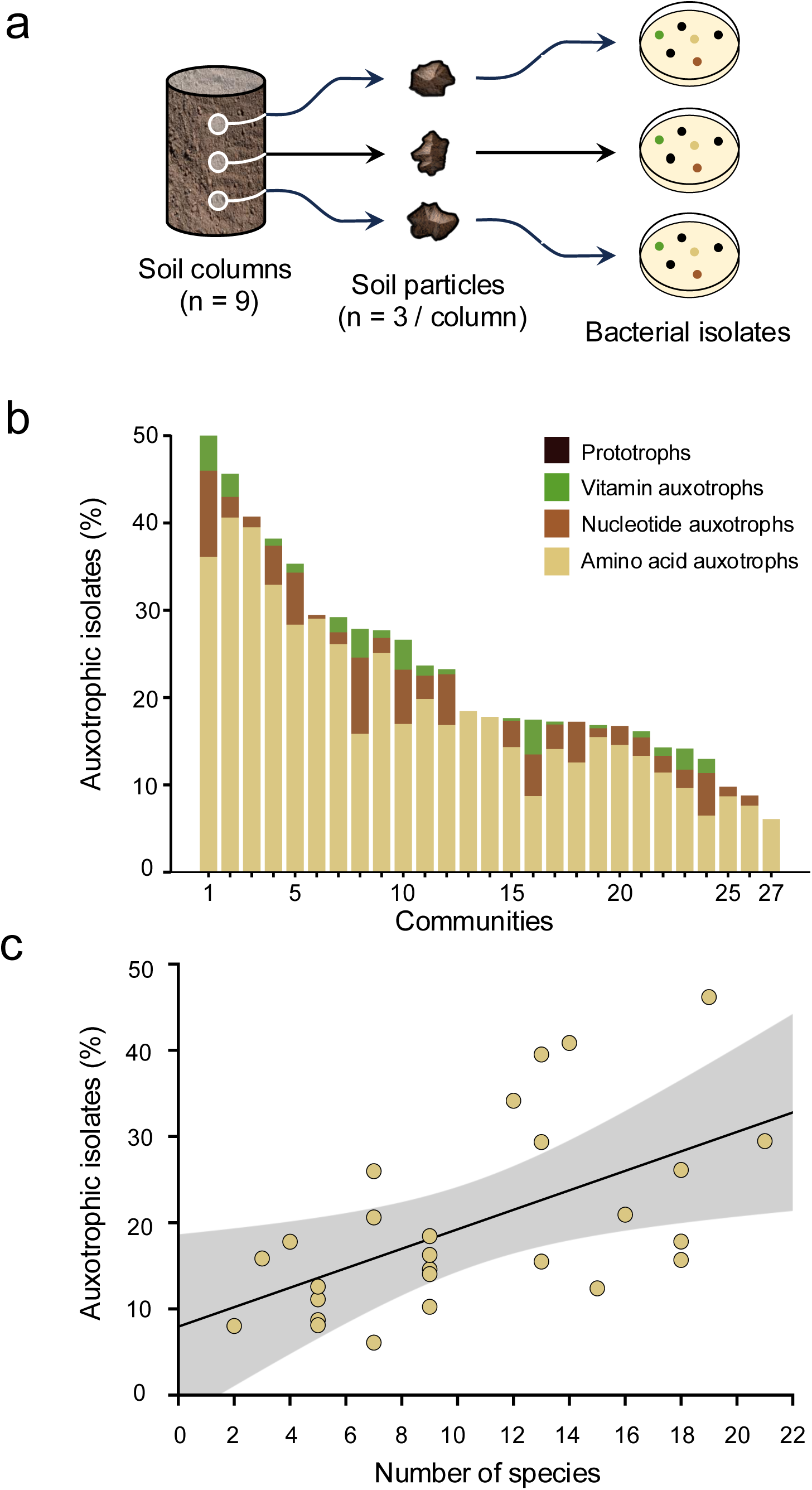
Auxotrophs are prevalent in soil microbial communities. (a) Nested experimental design to isolate 27 microbial communities from nine sites. A soil column of each site was collected and analysed. Of each column, three soil particles (1 mg each) were selected and used for bacterial isolation. (b) Proportion of auxotrophic isolates and type of metabolic auxotrophy of each community. (c) Relationship between the number of species per soil particle (i.e., species richness) and community-level proportion of auxotrophs detected for the corresponding community (Spearman correlation: (***ρ***) = 0.6, p = 9.7 x 10^-4^, n = 27).

Next, all single isolates and mixed clones were subjected to a plate-based assay in order to determine their ability to grow without supplementations (i.e., amino acids, vitamins, and nucleosides; Methods). The results revealed that between 10 -50% of all tested bacteria were auxotrophic for at least one tested supplement (Fig. 1b). Notably, a large number of strains were unable to grow in the absence of amino acids (i.e., 6 - 39%, median = 16%), while a smaller subset required vitamins (i.e., 0.4 - 10%, median = 2%) or nucleosides (i.e., 0.3 - 2%, median = 0.7%, Fig. 1b) to grow. Interestingly, communities from different soil particles showed a marked difference in their relative proportion of auxotrophic isolates (Fig. 1b). Taxonomically identifying 700 isolated auxotrophs by sequencing their 16S rRNA gene revealed a significant correlation between the number of species per soil particle (i.e., community richness) and the proportion of auxotrophs detected on the community-level (Fig. 1c). This finding suggests that more diverse microbial communities can support a higher number of auxotrophic isolates. Together, these results showed that amino acid auxotrophic strains are common in soil microbial communities.

### Isolated strains are commonly auxotrophic for multiple amino acids

The isolated amino acid auxotrophs were further characterised to determine which specific amino acid(s) each strain required for growth. For this, amino acid auxotrophs were inoculated on twenty different solid media, each containing a different mixture of nineteen amino acids (Methods). The vast majority of auxotrophs were unable to autonomously produce multiple amino acids (73%, Fig. 2a). While only 27% of the tested strains were auxotrophic for only one amino acid, most strains were auxotrophic for two (30%) or three amino acids (35%, Fig. 2a). Interestingly, 0.5% of all isolates could not grow on any of the single amino acid drop-out plates, yet showed good growth in the presence of all 20 amino acids (Fig. 2a).

**Fig 2.**
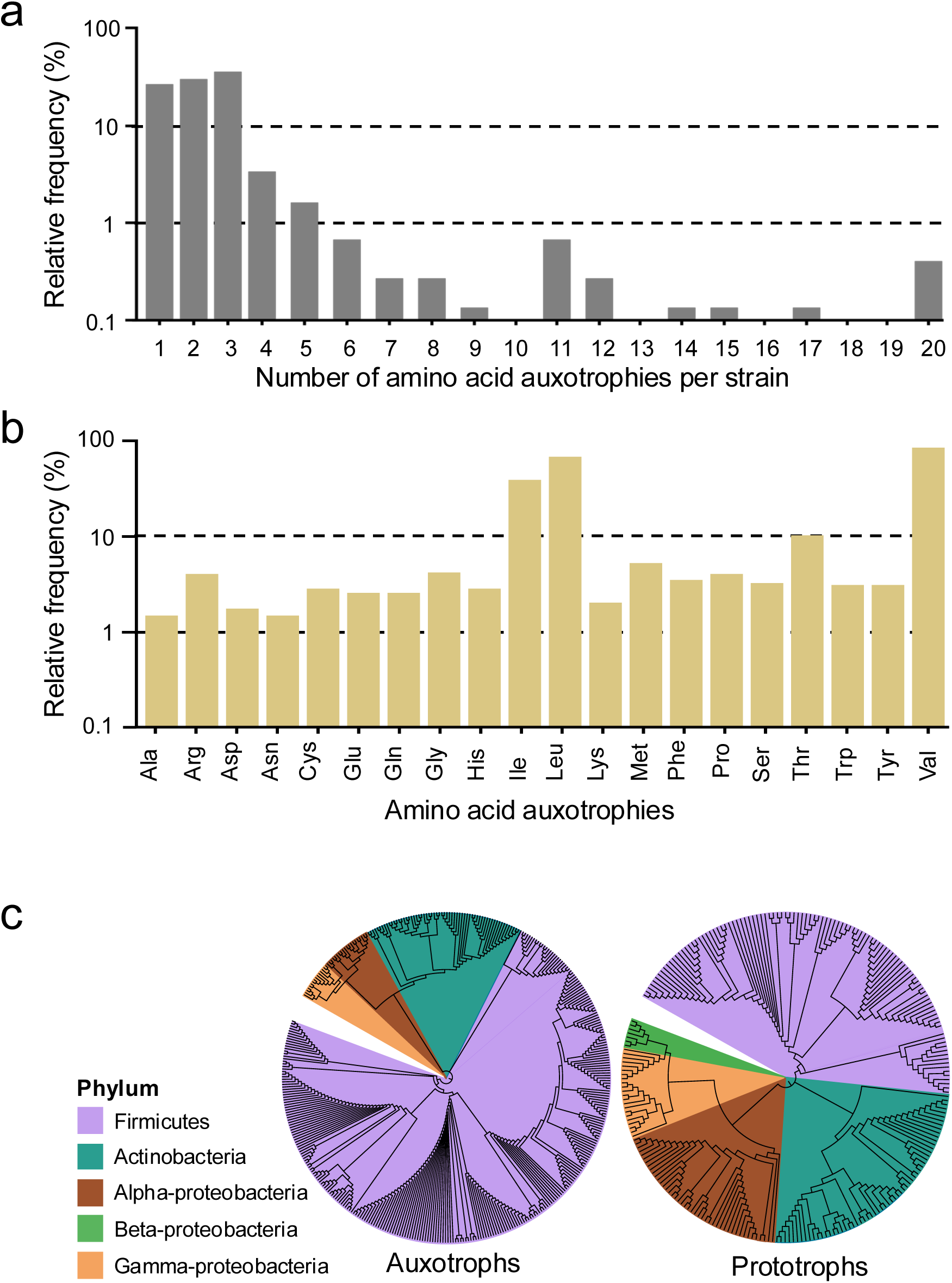
Soil isolates are frequently auxotrophic for multiple amino acids and commonly lack the ability to produce branched-chain amino acids. (**a**) Relative frequency of the number of amino acids auxotrophies per strain in the amino acids auxotrophs identified. **(b**) Relative frequencies of each amino acid auxotrophy in the screened set of amino acids auxotrophs. (**c**) Phylogenetic distribution of all isolated auxotrophs (n = 390) and representative prototrophs (n = 240) based on the sequencing of their 16S rRNA gene. The phylogenetic trees were generated using iTOL ^60^.

The most commonly observed auxotrophies were valine (59%), leucine (47%), and isoleucine (27%, Fig. 2b). In contrast, auxotrophies for asparagine (1%), alanine (1%), and aspartic acid (1%) were least common (Fig. 2b). All other auxotrophies occurred in a frequency of less than 4% of all analysed isolates. An exception was the auxotrophy for threonine that occurred in 7% of all auxotrophs (Fig. 2b).

Together, these results revealed that strains isolated from soil were frequently auxotrophic for multiple amino acids simultaneously and that auxotrophies for valine, leucine, and isoleucine were most commonly observed.

### Auxotrophs belong to various classes of indigenous soil inhabitants

16S rRNA gene sequencing was used to determine the phylogenetic affiliation of 390 amino acids auxotrophs as well as of 240 randomly chosen prototrophic strains. Both auxotrophs and prototrophs mainly belonged to three bacterial phyla: Firmicutes, Actinobacteria, and Proteobacteria (Fig. 2c). After excluding phylogenetically similar strains, auxotrophic and prototrophic isolates were found to share a similar phylogenetic distribution, with Firmicutes being most prevalent. Finally, sequencing metagenomes of the nine study sites (one sample per site) revealed a diversity that vastly exceeded the diversity of isolated strains (Supplementary Fig. 2).

### Comparing context-specific metabolic model predictions to experimental observations reveals an underestimated number of auxotrophies

Next, we questioned whether the genomic content of isolated strains or context-specific genome-scale metabolic models (GEMs) could explain the observed pattern of amino acid auxotrophies. To test this, we sequenced a subset of 64 isolates the 9 metagenomic samples from which they were cultured. These samples were processed using the metaGEM^14^ workflow to assemble genomes and reconstruct GEMs. A total of 102 genomes were generated with 70 SAGs (single amplified genomes) and 32 MAGs (metagenome assembled genomes). Curiously, four isolate samples yielded two high-quality genome assemblies per sample and one sample yielded three high-quality genomes (Supplementary Fig. 3a).

GEMs were used to predict amino acid auxotrophies based on flux balance analysis (FBA) simulations, which were then compared to experimental observations of amino acid auxotrophies (Fig. 3a, Supplementary Fig. 4). Although many auxotrophies were observed experimentally (Fig. 2a, b), the majority of assays resulted in prototrophic observations, thus creating a so-called imbalanced dataset for evaluation of auxotrophic predictions (i.e. 81.7% prototrophic versus 18.3% auxotrophic). In short, laboratory observations used for evaluation consisted of binary (true or false) auxotrophy data for 20 amino acids across 62 genomes (originating from 58 isolated samples), resulting in a total of 1,240 observations (Fig. 3a, b). Out of the 227 strain-amino acid auxotrophy combinations, only 17 true positives could be correctly predicted by GEMs, while a total of 1,013 true negatives were correctly identified. In addition, 210 false negatives were predicted (i.e. cases, where the models predict prototrophy, but an auxotrophy has been detected experimentally). The metabolic model predictions had a Spearman correlation (***ρ***) of 0.36 with *P* value = 0.004, suggesting that these GEM-based predictions are not able to fully account for the observed pattern of auxotrophies. Notably, no false positives were predicted, suggesting that the models likely underestimate rather than overestimate the true number of metabolic auxotrophies.

**Fig 3.**
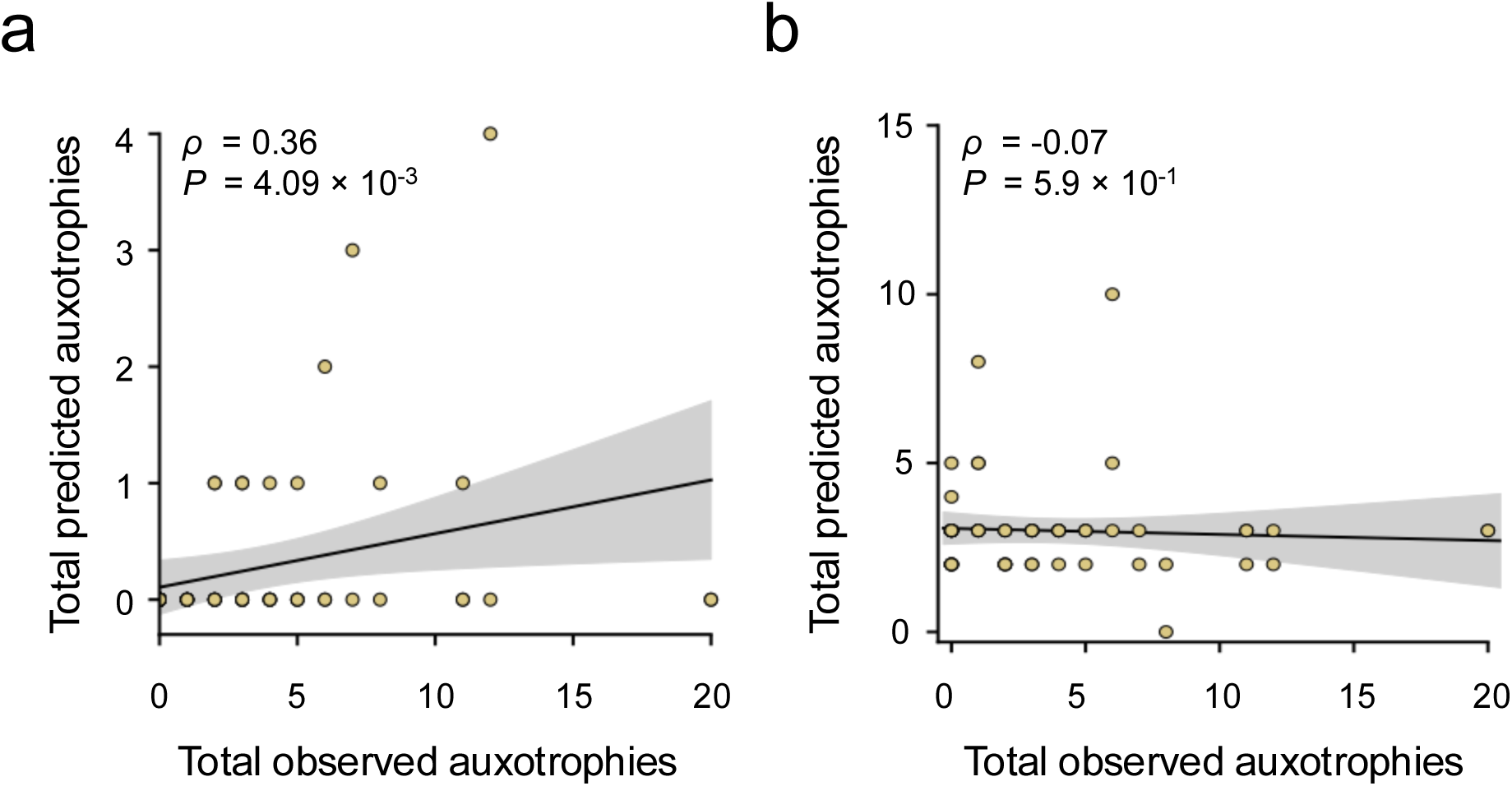
Metabolic modelling and analyses based on genome annotations fail to accurately predict experimentally observed auxotrophies. Comparison between the predicted number of amino acid auxotrophies based on (**a**) genome-scale metabolic modelling and (**b**) genome annotations, respectively, and the number of observed amino acid auxotrophies across sequenced isolates. Scatter plots show regression lines and shaded areas correspond to 95% confidence intervals

As expected, genomic annotations of biosynthesis pathway by themselves performed worse than GEMs (Supplementary Fig. 4). To assess the predictive performance of biosynthetic gene annotations, we used KEMET^15^ and eggNOG^16^ to estimate pathway completeness for 14 amino acid biosynthesis routes and evaluated them against the experimental observations (Fig. 3b). Genomes were considered prototrophic for a given amino acid if they contained at least one complete biosynthesis route; otherwise they were considered to be auxotrophic. The genomic content was able to correctly identify 33 true positives and 512 true negatives, while suffering from 141 false negatives and 149 false positives. This model had ***ρ*** = -0.07 with a *P* value of 0.59, indicating that the gene annotation-based predictions cannot fully capture the observed pattern of auxotrophies, largely due false positives (Fig. 3b, Supplementary Fig. 4).

Together, these results clearly showed that the approaches used to predict amino acid auxotrophies were unable to account for the experimentally observed metabolite auxotrophies.

### Genomic mechanisms driving the evolution of amino acid auxotrophies

Taking advantage of the sequenced microbial genomes, we aimed to identify the molecular mechanisms that caused the evolution of auxotrophic phenotypes among soil bacteria. To address this, the assembled genomes were analysed to determine the role of four possible drivers: i) loss of chromosomal biosynthetic genes, ii) accumulation of insertion sequences (ISs), iii) genomic changes associated with mobile genetic elements (i.e., plasmids and viruses), and iv) metabolic or transcriptional regulation (Supplementary note 1). When comparing auxotrophic and prototrophic genomes, there was no clear difference in the number of plasmid or virus-encoded genes (Supplementary Fig. 5a, b, Supplementary note 2). However, there were significant differences between auxotrophs and prototrophs in terms of the number of ISs found in overall annotated genes, as well as genes annotated with amino acid metabolism (Supplementary Fig. 5c, d).

#### (i) Depletion of specific genes in genomes with auxotrophic behaviour

To verify whether gene loss drove the emergence of amino acid auxotrophies, we annotated our genomes with the orthology-based functional annotation tool eggNOG mapper^16^ and the associated database^17^. There was no significant difference in genome size between auxotrophs and prototrophs (Supplementary Fig. 6a). On average, we identified a total of 4,140 annotated genes per genome (SD = 1,444) and this number that did not differ significantly between genomes of auxotrophic versus prototrophic isolates (Supplementary Fig. 6b). However, we did observe significant differences in genomic coding density, where auxotrophs (median = 934, IQR = 56.9, n = 9) had a significantly greater number of annotated genes per genome megabasepair (Mann-Whitney U test: *P* = 0.008, W = 106) compared to prototrophs (median = 880, IQR = 33.4, n = 53), which is one of the five traits commonly associated with genome streamlining^18^ (Fig. 4a). Similarly, there was a significant enrichment in the number of annotated genes per megabasepair (Mann-Whitney U test: *P* = 0.003, W = 687) in metagenome assembled genomes (N = 32, median = 1001, IQR = 222) compared to isolates (N = 68, median = 928, IQR = 65.5).

**Fig. 4.**
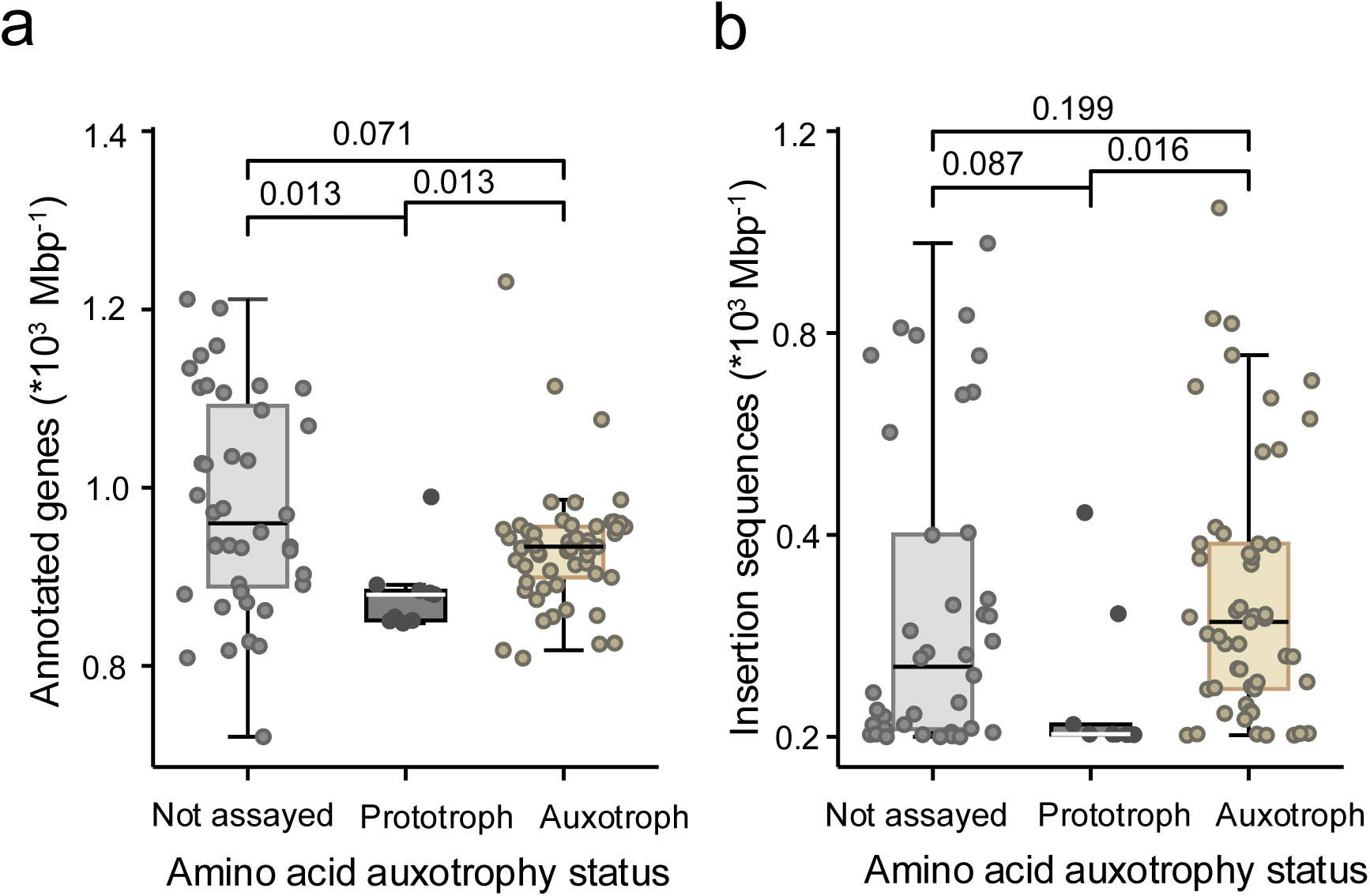
Genomic coding density and number of insertion sequences is increased in genomes of auxotrophic strains. Number of (**a**) annotated genes and (**b)** insertion sequences per genome megabasepair across metagenomic (not assayed), prototrophic, and auxotrophic isolates, respectively. In total, 102 isolated and metagenomically assembled genomes were analysed. Boxplots show the median (middle line), lower (25%), and upper (75%) quartiles, and whiskers represent the 1.5 interquartile range. Benjamini-Hochberg corrected P-values of Mann-Whitney U tests are shown.

Out of the 12,827 annotated genes identified across genomes, 260 showed a copy number that differed significantly between genomes of auxotrophic and prototrophic phenotypes (Benjamini-Hochberg corrected Mann-Whitney U test: *P* < 0.05). Within these genes, 30 were enriched and 230 were depleted in genomes of auxotrophic relative to prototrophic isolates (Supplementary note 3). A principal component analysis (PCA) of gene copy number across genomes did not show good separation between genomes with auxotrophic and prototrophic behaviour (Supplementary Fig. S7).

#### (ii) Insertion sequences are enriched in genomes of auxotrophic isolates

ISs and mobile genetic elements have recently gained attention, due to their role in the evolution of metabolic auxotrophies and the streamlining of genomes^19,20^. For example, researchers have demonstrated that enrichment of ISs in *Shigella* species resulted in genomes of reduced size and a concomitant loss of metabolic capabilities^21^. This mechanism provides a powerful explanation for the phenomenon of biosynthetic gene loss, whereby ISs accumulate to the point of knocking out metabolic functions and, ultimately, leading to gene loss.

The hypothesis that ISs are associated with the evolution of amino acid auxotrophies was tested by using the ISfinder database^22^ (i.e., a set of 8,433 prokaryotic ISs) in combination with the DIAMOND^23^ sequence aligner to query their presence and distribution across genomes. The median number of ISs per genome megabasepair was 165 (IQR = 358), with a median of 46.8% sequence identity similarity per hit (IQR = 11.5%), and a median of 100 bitscore per hit (IQR = 123). Notably, we found that the number of ISs per megabasepair was significantly higher in genomes that derived from auxotrophic isolates (median = 227, IQR = 288, n = 53) as compared to their number in prototrophic genomes (median = 5.1, IQR = 19.8, n = 9) (Benjamini-Hochberg corrected Mann-Whitney U test: *P* = 0.016, W = 99, n = 62) (Fig. 4b, Supplementary note 4).

In sum, the observed enrichment of ISs in genomes of auxotrophic relative to prototrophic isolates supports the hypothesis that ISs likely play an important role in the emergence of metabolic auxotrophies.

### Amino acid cross-feeding stabilizes the growth of amino acid auxotrophs

Given the observed ubiquity of amino acid auxotrophies in soil-derived bacteria, we sought to understand the mechanism that stabilizes these loss-of-biosynthetic-function mutants in the corresponding microbial communities. One possible explanation is the exchange of metabolites with other members of the local community.

To determine whether a diffusion-based exchange of amino acids can explain the persistence of auxotrophic isolates, a bacterial community isolated from one soil particle was chosen (i.e., number 7 in Fig 1b). This community was selected based on its high phylogenetic diversity of both auxotrophic and prototrophic isolates. In total, 19 auxotrophic and 22 prototrophic strains were cocultured in all possible pairwise combinations involving either pairs of two auxotrophs (n = 361) or auxotroph-prototroph combinations (n = 418). Each combination was tested in four biological replicates.

The results revealed that 31% of auxotrophic genotypes tested displayed detectable growth when cocultured with another auxotroph (Fig. 5a), while 84% could grow when cocultured with prototrophic donors (Fig. 5a). Strikingly, the growth of auxotrophic recipients was supported by at least 1 other auxotroph and 7 prototrophic strains tested, suggesting that cross-feeding of essential metabolites is potentially very common in these communities.

**Fig 5.**
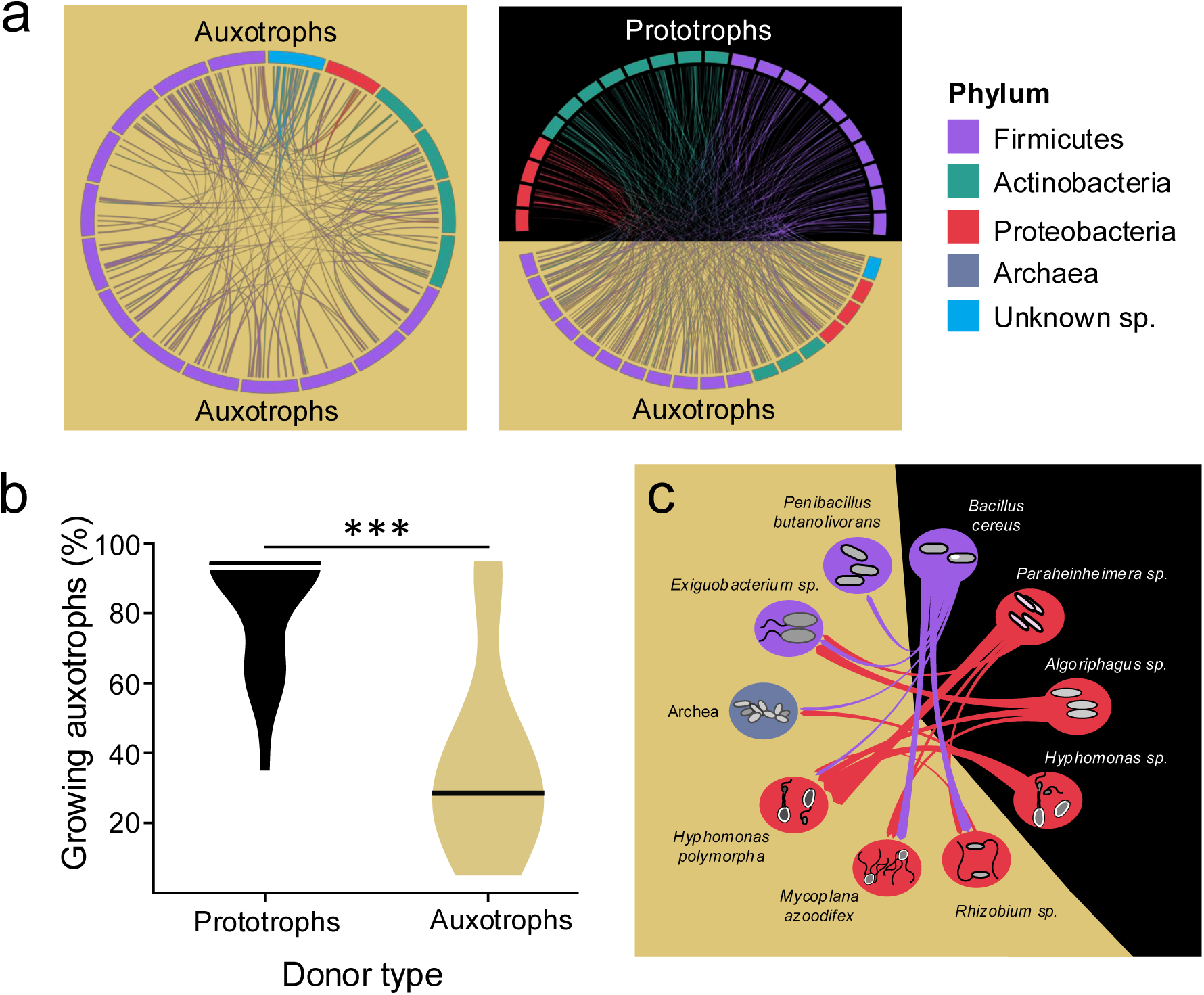
Cross-feeding with co-occurring strains stabilizes amino acid auxotrophs. (**a**) Interaction network of experimentally determined cross-feeding interactions in pairwise cocultures of two auxotrophs (left) and a prototroph and an auxotroph (right). All used strains have been isolated from the same soil particle (i.e., 19 auxotrophic and 22 prototrophic strains). The colour of nodes indicates the taxonomic identity of focal strains. Edges indicate the ability of auxotrophs to grow in coculture with the other strain and its colour indicates the taxonomic identity of the metabolite donor. (**b**) Growth of auxotrophic isolates was significantly enhanced in cocultures with prototrophs relative to cocultures with auxotrophic strains. Asterisks (***) show the results of a Mann-Whitney-U test: *P*-value = 0.000192, U = 350, n (prototrophs) = 22, n (auxotrophs) = 19. (**c**) Predicted metabolic interactions between auxotrophic and prototrophic assembled genomes from the metagenome sequencing of the same community as analysed before (a,b). The type of exchanged metabolites is shown in Supplementary Fig. S10.

In general, prototrophic donors were significantly more likely to support the growth of auxotrophic than was the case for auxotrophic donors (Fig. 5b). Interestingly, the ability of auxotrophs to grow in coculture with different auxotrophic or prototrophic isolates varied widely among the analysed cocultures (auxotrophic donors: median = 30%, IQR = 22.5; prototrophic donors: median = 95%, IQR = 30). This pattern was likely due to the number of amino acids a given auxotrophic strain required for growth: the more amino acids it could not produce autonomously, the less likely it was to grow in coculture with a metabolite donor. A statistical analysis of this interpretation revealed indeed a negative significant relationship for the case of prototroph-auxotroph cocultures (Spearman correlation: (***ρ***) = - 0.66, P = 2.2×10 ^- 5^, n = 22, Supplementary Fig. 8). However, no statistically significant relationship between the amount of amino acids prototrophic isolates released and the growth auxotrophic isolates showed in coculture could be detected (Supplementary Fig. 9 and Supplementary note 5).

These observations raised the question whether multi-auxotrophic strains are embedded in a cross-feeding network that includes more than one partner. To address this, we investigated the GEMs that were associated with the focal soil community (i.e., number 7 in Fig 1b). Specifically, metabolic interactions for this community were predicted using SMETANA^24^. To improve visualization, we focused our analysis on an exchange of vitamins and amino acids (Fig. 5c, Supplementary Fig. 10) and excluded other compounds (i.e., ions, trace metals, and carbon sources). Ten additional genomes were assembled from metagenomic samples, which could not be isolated from the corresponding bacterial community. The inferred auxotrophs were found to be multi-auxotrophic for both amino acids and vitamins (Supplementary Fig. 10) and their growth was likely facilitated by cross-feeding with more than one prototrophic genotype (Fig. 5c, Supplementary Fig. 10). The only exception was a thiamine auxotrophic *Peribacillus butanolivorans* strain, which likely depended on *Bacillus cereus* to grow. Interestingly, the model also revealed potential cross-feeding between auxotrophic strains. For example, *Rhizobium sp*. (auxotrophic for thiamine and threonine) was predicted to provide homoserine to *Hyphomonas sp*., which is likely auxotrophic for four different metabolites. Together, these results support the hypothesis that multi-auxotrophic strains thrive by cross-feeding with multiple partners.

The mixture of metabolites that is produced by several, genetically different strains is likely more chemically diverse than the blend produced by a single strain. Consequently, the probability that an auxotrophic recipient can grow should be higher when interacting with multiple strains simultaneously as compared to cocultures with only one partner. In addition, multipartite interactions also increase the likelihood of complementary auxotrophies, thus further enhancing the potential for metabolic cross-feeding.

To investigate the potential for metabolic complementarity among auxotrophic bacteria, we analysed the complete set of 746 auxotrophic strains that derived from 27 communities (Fig 1b). Specifically, we assessed whether groups of two, three, four, or five strains could collectively produce the complete set of 20 amino acids, thus ensuring that each auxotrophic dependency was complemented by at least one member of the group. This analysis revealed a clear trend, in which increasing group size was associated with a higher likelihood of complete auxotrophic complementarity, suggesting that the potential for metabolic support through cross-feeding interactions is increased in larger consortia (Fig 6a).

**Fig 6.**
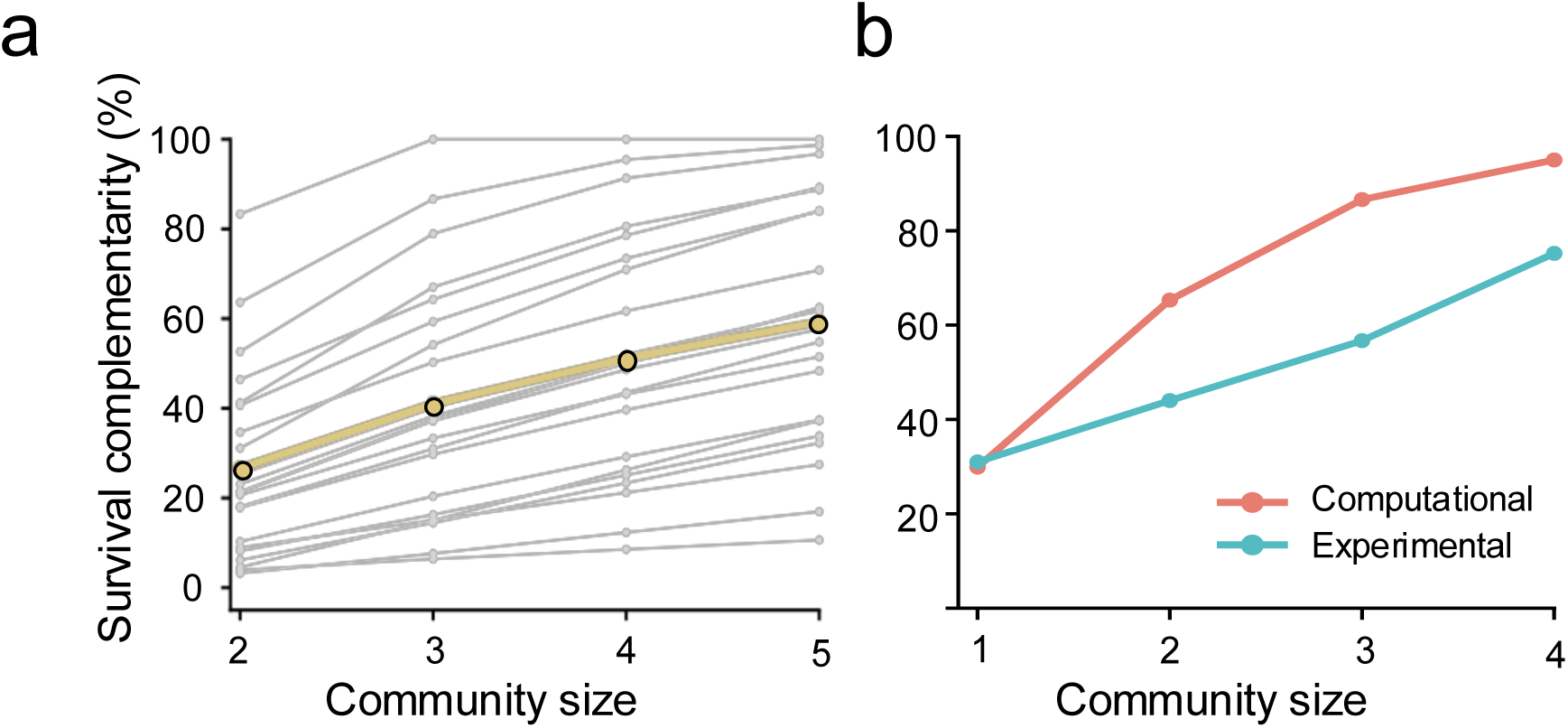
Metabolic complementarity within an auxotrophic community increases with an increasing community size. (**a**) Complementarity of amino acid auxotrophies among groups of co-occurring auxotrophs across 27 soil communities. Gray lines represent complementarity for each of 27 community as group size increases from pairs to 5-way interactions, while the beige line shows the mean complementarity across all communities. Communities with consistent zero complementarity across group sizes were excluded from the plot (n = 5). (**b**) Degree of complementarity of lab-observed auxotrophies in pairwise and higher-order combinations of 58 isolates, based on computational analysis (red) and empirical testing using 12 isolates (blue).

Next we sought to experimentally verify the observed pattern. For this, we focussed on all 58 sequenced isolates of the focal community (i.e., number 7 in Fig 1b) that included both auxotrophic and prototrophic phenotypes. First, we subjected these strains to the same computational analysis as before. In this way, we aimed to obtain a baseline against which experimental results can be compared. Of the focal strains, we analysed all non-redundant 58 monocultures and 3,305 pairwise combinations, as well as 3,356 randomly assembled combinations consisting of three or four isolates. As before, auxotrophies were assumed to be complemented if at least one group member was prototrophic for the focal amino acid. In line with previous observations, the results of this computational analysis revealed a complementarity of 29.3%, 65.1%, 86.6%, and 95.1% across pairwise, three-way, and four-way combinations, respectively (Fig 6b).

Finally, we validated these obtained results experimentally by analysing cultures consisting of one, two, three, or four isolates from a total pool of 16 strains. All combinations were randomly chosen, and - to avoid combinatorial explosion – limited to 120 cocultures for each of the pairwise, three-way, and four-way combinations. Cocultures were inoculated on minimal media agar plates without supplementation (i.e., amino acids, vitamins, and nucleosides). Under these conditions, growth indicated a complete resolution of all auxotrophies. Across pairwise, three-way, and four-way combinations, we observed growth in 44.2%, 56.7%, and 75.0% of tested communities and populations, respectively (Fig. 6b). Our experimental data therefore corroborated the results of the abovementioned computational analyses (Fig. 6b). Thus, both modelling and experimental results support the hypothesis that increasing the number of genotypes in a microbial community enhances the likelihood for a metabolic complementarity among co-occurring genotypes.

## Discussion

An obligate exchange of essential metabolites has been suggested to be one of the main reasons for the frequently observed difficulty to cultivate environmental bacteria under laboratory conditions^3^. Here, we experimentally investigated the prevalence of metabolically obligate-dependent isolates in soil bacterial communities, analysed their genomic basis, and tested their ability to establish cross-feeding interaction with other community members. To this end, we experimentally screened > 6,000 bacterial strains that have been isolated from 27 soil microbial communities. We found that up to 50% of bacterial strains from a given community showed an auxotrophic phenotype, with amino acid auxotrophies being most commonly observed. Furthermore, assembly-based bioinformatic analysis of metagenomes and isolates allowed us to reconstruct full genomes as well as the corresponding GEMs. Genomic inspection identified (i) an increased coding density, (ii) a depletion of 230 genes, and an enrichment of ISs in auxotrophic genomes compared to genomes of prototrophic strains. Combining systematic coculture experiments with computational analyses revealed that the long-term persistence of auxotrophic isolates in these environments is likely due to a diffusion-based exchange of amino acids between different members of the local community.

Our results reveal that multi-auxotrophic isolates are abundant in soil microbial communities (>70% of the total isolated auxotrophs). Interestingly, auxotrophies for branched-chain amino acids (BCAAs, i.e., valine, leucine, and isoleucine) were most commonly observed. Several reasons can explain this pattern. First, valine is the amino acid that is released the most from bacteria cells^25^, while leucine is most abundant in proteins across living cells^26^. Thus, leakage and degradation of dead cells will likely result in an accumulation of valine and leucine in local environments, which in turn should promote the evolution of the corresponding auxotrophies. Second, BCAAs share genes in their biosynthetic pathways^27^. Consequently, a mutation or loss of one gene is likely to also affect the production of the other two metabolites. Third, acquiring auxotrophies for BCAAs increases a cell’s resistance to antibiotics. This is because auxotrophic mutants of BCAAs have reduced level of branched-chain fatty acids in their membrane phospholipids^28^. The resulting decrease in fluidity^28^ and permeability^29^ of the outer membrane prevents various antibiotics from entering the cell. Thus, both the availability of these amino acids and the interplay with other ecological factors are likely to shape the occurrence of specific amino acid auxotrophies.

Previous studies have explored the abundance of metabolic auxotrophies in soil bacterial communities^30,31^. The results of these analyses, which mainly relied on culture-independent approaches, corroborated our finding that auxotrophs are common in these environments^30,32^. However, some studies yielded contradictory patterns. For example, a recent large-scale screening of 26,000 bacterial isolates^31^ combined a 16S-based sequence analysis with metabolic modelling to predict amino acid auxotrophies. The authors of this study concluded that amino acid auxotrophies were generally rare in soil environments^31^. Interestingly, our own attempts to predict experimentally observed auxotrophic phenotypes by analysing the reconstructed metabolic networks of sequenced isolates also underestimated the number of experimentally confirmed auxotrophies drastically (Fig. 3a). Several reasons might explain the observed discrepancy. First, metabolic auxotrophies may not only be caused by a loss of a biosynthetic gene, but also by as-of-yet unknown pleiotropic mutations, through epigenetic mechanisms, or mobile genetic elements (MGEs)^20^. For example, regulatory changes can cause auxotrophic phenotype, even though the biosynthetic machinery to produce amino acids remains intact^33,34^. Moreover, ISs can also affect neighbouring genes^35^. Consequently, genetic screens that determine whether or not the central biosynthetic genes are fully present in the genome^15^ are likely to underestimate the true number of metabolic auxotrophies. Second, a large number of genes across natural isolates and yet-uncultured strains are not functionally annotated^36^. Thus, existing links to the strain’s amino acid metabolism remain undetected. Indeed, researchers have recently characterised a previously unknown sequence space across metagenomic samples^37^. The results of this analysis suggest that some of the most widely spread novel protein families were associated with amino acid metabolism and transport in soil microbiomes (Table S2^37^, e.g. F000025 & F000036). Third, many “known” gene annotations that are present in databases may be inaccurate or incorrect^38^, thus leading to discrepancies between expected and observed auxotrophs.

Overall, our study demonstrates that cultivation-based approaches are essential to accurately determine the number of auxotrophs in natural microbial communities. Given that the majority of environmental bacterial strains resist cultivation under laboratory conditions, novel approaches that allow the isolation of whole metabolically-dependent consortia should be used^39–41^.

Based on our sequencing data, we conclude that an accumulation of ISs in auxotrophic genomes is a likely cause for the evolution of auxotrophic phenotypes. From an evolutionary point of view, this mechanism offers a mechanistic explanation of the emergence of auxotrophies without entirely losing the corresponding biosynthetic machinery. Depending on the local availability of this compound, the auxotrophic cell will either be able to grow or is doomed to die because of starvation. However, environmental conditions are subject to continuous and rapid change. Under these conditions, ISs that disrupt^42^ or regulate genes^35^, serve as a bet-hedging strategy that provides two different advantages. First, the phenotypic consequences of IS-based genomic modifications are potentially reversible^43^. This ensures the survival of a given population of cells. Second, IS-mediated mechanisms will increase the variation of a local population with regard to its metabolic capabilities and requirements, thus enhancing its ability to cope with fluctuating environmental conditions. In the long run, this process will likely result in streamlined genomes as expendable genes are eventually lost from the genome due to an accumulation of ISs or IS-mediated genomic recombination^21^. Viewed through this lens, our study identified ISs and the resultant ecological dynamics as a likely driver for the commonly observed unculturability of environmental bacteria.

Taken together, our work shows that auxotrophic bacteria are common in natural microbial communities and demonstrate that they are stabilized by a cross-feeding of metabolites with other co-occurring strains. In addition, our findings imply that bacteria are likely to exist within complex networks, within which they simultaneously interact with many other strains or species. Our study therefore identifies obligate cross-feeding of essential metabolites as a key mechanism to account for the frequently observed difficulty in cultivating environmental bacteria under laboratory conditions.

## Material and methods

### Culture conditions and general methods

In all experiments, minimal medium for *Azospirillum brasiliense* (MMAB) ^13^ that contained fructose (5 g l^-1^) as sole carbon source has been used as the main medium. Depending on the experiment, this medium was supplemented with all twenty proteinogenic amino acids (200 µM each), five vitamins (i.e., riboflavin (B2), cobalamin (B12), biotin (B7), thiamine (B1), and pyridoxine (B6); 1 µM each), as well as four nucleobases (i.e., cytosine, adenine, uracil, and thymine; 100 µM each). Agar plates used to isolate bacteria from soil samples have been additionally supplemented with the fungicide Nystatin (100 µg ml^-1^).

To prepare precultures, the respective strains were streaked from glycerol stocks on agar plates and incubated for five days at room temperature. A single colony of the isolate was then transferred into a 96 deep-well plate containing 1 ml liquid medium and incubated under shaking conditions (200 rpm) for 48 hours at 30 °C.

### Collection of soil samples

Nine soil samples were collected, which were all located around Jena in Thuringia, Germany (Fig. S1). After removing the surface layer of soil (i.e., top 5 cm), a T-sampler was used to collect a soil column of 5 cm in diameter and 5 cm in length. Sampled columns were transported to the laboratory without disrupting their structure. Under sterile conditions, columns were manually dissected. Three soil particles (volume: ca. 1 mm^3^, weight: 1 mg), which were 2.5 cm apart, were selected for bacterial isolation. This nested design resulted in the sampling of a total of 27 different soil microbial communities (Fig. 1a).

### Bacterial isolation and purification

Each selected soil particle was suspended in 500 µl of saline solution (0.9% NaCl) and shaken for 30 minutes at 200 rpm. 100 µl of this solution was then transferred into 400 µl of saline solution and vortexed for two minutes. For bacterial isolation, 100 µl aliquots of each dilution were spread-plated on MMAB agar plates of the above-described medium (i.e., three replicates per dilution) and incubated at room temperature for five days. All growing clones (n = 6,931) were purified at least twice by sub-streaking to obtain pure colonies and stored as glycerol stocks (1 ml liquid culture + 1 ml glycerol (50%)) at -80 °C until further use.

### Screening for metabolic auxotrophies

Precultures of all isolated clones were prepared as described above. 2 µl of growing precultures were inoculated on five different agar plates using a 96-pin replicator (Boekel Scientific 140500 Microplate Replicator). The five screening media consisted of MMAB as a base, yet differed in the type of supplements added. The first medium contained amino acids, vitamins, and nucleobases, which should allow both prototrophs and auxotrophs to grow (i.e., positive control). The second medium consisted of MMAB without any supplements added. On these plates, only prototrophic bacteria should be able to grow (i.e., negative control). The third medium contained MMAB that has been supplemented with vitamins and nucleobases. Amino acid auxotrophic strains should not be able to grow on this media. The fourth medium contained vitamins and amino acids, thus preventing nucleobase auxotrophic strains from growing. The final medium contained amino acids and nucleobases, on which genotypes auxotrophic for vitamins should not be able to grow. Inoculated plates were incubated at room temperature for four days. Subsequently, the growth pattern of each isolate across the five different media was analysed. An isolate was considered auxotrophic if it could not grow on unsupplemented agar plates (i.e., negative control), but any of the other plates. The type of auxotrophy was determined by observing a lack of growth on plates, from which a specific group of metabolites (i.e., amino acids, vitamins, or nucleobases) has been omitted.

### Determining the profile of amino acid auxotrophies

To determine for which amino acid(s) the isolated strains were auxotrophic for, all identified amino acid auxotrophic strains (i.e., 1,494 clones) were revived from frozen stocks and precultured as described above. Afterward, isolates were inoculated on 22 different media using the 96-pin replicator. All media consisted of MMAB, to which a mixture of five vitamins has been added. This time, however, nucleotides were not added to the medium, because of an observed growth-inhibiting effect on many of the isolated strains. The 22 different media contained (i) no amino acids (i.e., negative control), (ii) all 20 proteinogenic amino acids simultaneously (i.e., positive control), or (iii) a mixture of nineteen amino acids, which differed in the identity of one amino acid that has been omitted (i.e., 20 different drop-out media). All tested isolates were expected to be unable to grow on unsupplemented media. If observed otherwise, the respective isolate was re-classified as ‘prototrophic’ and removed from further analysis. The profile of amino acid auxotrophies of each isolate was determined by comparing its growth pattern on all screening media. An isolate was considered auxotrophic for a specific amino acid if it could not grow on a medium that lacked the corresponding metabolite. For example, an isolate that could not grow on a medium supplemented with all amino acids except leucine was considered to be auxotrophic for leucine.

### Molecular phylogenetic analysis

To determine the phylogenetic identity of the isolated strains, 1,000 clones were subjected to 16S rRNA gene sequencing. The isolates selected for sequencing included 600 amino acids auxotrophic strains and 400 randomly chosen prototrophic genotypes. Isolates were revived and precultured as described before. Next, a 20 µl PCR reaction was set up by directly using 1 µl of the growing preculture as DNA template to which JumpStart™ REDTaq ReadyMix (Sigma Aldrich) as well as the general eubacterial primers FP (AGAGTTTGATCCTGGCTCAG) and RP (ACGGCTACCTTGTTACGACT) have been added. The PCR program used was: 3 min at 94 °C, 32 cycles of 40 s at 94 °C, 60 s at 65 °C, 60 s at 72 °C, and a final extension step of 4 min at 72 °C. PCR products were checked on agarose gel, purified using Shrimp alkaline phosphatase (Illustra, Fischer scientific), and sequenced at the Department of Entomology of the Max-Planck-Institute for Chemical Ecology in Jena, Germany. Sequencing was performed using an ABI 3730XL capillary DNA sequencer (Applied Biosystems, USA). The quality of retrieved sequences was verified manually. Both ends of the resulting sequences were trimmed and contigs were assembled. The resulting sequences of 16S rRNA genes, 1-1.2 kb in most cases were submitted to NCBI BLASTn for taxonomic identification and focal strains were assigned to the most closely related species using a minimum query coverage of 98% and minimum identity value of 99%.

### Pairwise coculture experiment

To test whether other strains can promote the growth of amino acid auxotrophic genotypes, a large-scale pairwise coculture experiment was performed following a previously published protocol^44^. For this, 18 auxotrophic and 23 prototrophic strains were selected that all had been isolated from the same soil particle and thus belonged to the same local community. The focal strains belonged to three bacterial phyla: Actinobacteria, Firmicutes, and Proteobacteria (Supplementary Table 1). A preculture of each isolate was prepared by transferring a few colonies into 3 ml of liquid medium, which was then incubated at 30 °C under shaking conditions (200 rpm) until each isolate had reached its exponential growth phase. Next, growing cultures were poured into 15 ml of fresh liquid medium, incubated as before, and, after reaching the exponential growth phase, transferred into 50 ml of liquid medium. Finally, cells were centrifuged, the supernatant discarded, and cells resuspended in 30 ml MMAB, which lacked a carbon source, vitamins, and amino acids. Next, the optical density (OD) of the prepared suspensions was measured at 600 nm using a FilterMax F5 multi-mode microplate reader (Molecular Devices) and adjusted to 0.1. Each strain was then mixed with melted MMAB agar (45 ml) that contained fructose as the sole carbon source, but lacked additional media supplements. The resulting cultures, which showed a final optical density of 0.01, were poured into square plates and allowed to solidify for two hours (hereafter referred to as ’donor’). Using a 96-pin replicator, aliquots of auxotrophic strains (OD_600nm_: 0.05) were then inoculated (i) on top of each donor strain, (ii) on a plate that contained amino acids and vitamins, yet lacked a donor strain (i.e., positive control), and (iii) on a plate that contained pure MMAB, yet lacked media supplements and a donor strain (i.e., negative control). All interaction and control plates were incubated at room temperature for five days and then imaged using the BIORAD GelDoc Go Imaging system (Biorad Laboratories GmBH). An image-processing model was developed to quantify the growth of colonies. The developed machine-learning codes can be accessed at https://github.com/franciscozorrilla/soil_auxo.

### DNA extraction

To identify the genetic basis of the observed metabolic auxotrophies, the genomes of 60 bacterial soil isolates were fully sequenced. Isolates were selected to cover the observed phylogenetic diversity and to represent different profiles of amino acid auxotrophies. In addition, two prototrophic genotypes have been included as a reference for complete amino acid biosynthetic pathways. All isolates were revived from glycerol stocks. For every strain, two liquid cultures (7 ml each) were inoculated and incubated for 48 hours at 30 °C under shaking conditions (200 rpm). Cells from both replicates were pooled, collected by centrifugation, and stored at -20°C until further use.

The genomic DNA was extracted following the protocol of Kowalczyk *et al.* ^45^ and later modified by Blasche *et al.* (2021). For this, frozen cells were transferred into 2 ml tubes, suspended in 600 μl TES buffer (25 mM Tris, 10 mM EDTA, 50 mM sucrose) with 20 mg ml^-1^ lysozyme (chicken egg white, Sigma Aldrich), and incubated at 37 °C for 30 min ^44^. Then, 0.5 g of glass beads (size: 100-250 µm) were added and samples were bead-beaten three times for 30 seconds with one minute of cooling interval in-between at 4 m s^-1^ using a FastPrep®-24 5G (MP, Germany). Next, 150 μl of 20 % SDS was added, tubes were incubated for 5 minutes at room temperature, and centrifuged at 14,000 rpm for 2 minutes. The resulting supernatant was transferred into a fresh tube, mixed with 10 μl Proteinase K (20 mg ml^-1^), and incubated for 30 min at 37 °C. For protein precipitation, 200 μl potassium acetate (5 M) was added to samples. After that tubes were incubated for 15 minutes on ice and centrifuged for 15 min at 14,000 rpm at 4 °C. The clear supernatant was subjected to phenol-chloroform extraction by adding a mixture of Phenol/Chloroform/Isoamyl alcohol (ROTI®) in a 1:1:1 ratio. The resulting solution was gently mixed and centrifuged for 15 min at 14,000 rpm at room temperature. Then, the aqueous layer was transferred into a fresh tube. The extraction step was repeated twice. Finally, DNA was precipitated by adding cold isopropanol (-20 °C) corresponding to twice the volume of the transferred aqueous layer. The resulting tubes were incubated for 2 hours at -20 °C. The quality of the isolated DNA was verified by agarose gel electrophoresis and its concentration was determined using a UV/VIS-spectrophotometer (Nanodrop One, Thermo Scientific). Samples were sent for sequencing to the gene core facility at EMBL, Heidelberg, Germany.

### Soil metagenome extractions

One gram of soil from each site (Supplementary Fig. 1) was subjected to metagenome isolation using the DNeasy PowerSoil Pro Kit (Qiagen) following the manufacturer’s protocol.

The quality of isolated metagenomes was verified on an agarose gel. Samples were sent for sequencing to the gene core facility at EMBL, Heidelberg, Germany.

### Sequencing and library preparation

Sequencing libraries were constructed following the protocol NEBNext Ultra II DNA library Prep Kit for Illumina (New England Biolabs, Inc.). Briefly, 50 ng of each DNA sample was fragmented by sonication (Sonorex RK102 Transistor, Bandelin GmbH), followed by library preparation according to the manufacturer’s protocol of NEBNext Ultra II DNA Library Prep Kit for Illumina, with 8 cycles of PCR to amplify the adaptor-ligated fragments (Biometra TAdvanced, Analytik Jena GmbH). Size selection and purification of the finished libraries were performed with a 0.8x SPRI bead (Beckman Coulter) cleanup.

An initial set of 9 isolates were deep-sequenced on a HiSeq 4000 flowcell with chemistry for 300 cycles and a 150 bp paired end read mode, with on average 38 million reads per sample. Samples from metagenomics experiments were pooled in a 5x scheme with on average 30 million reads per sample. Pooled samples were loaded onto a single NextSeq 500 High flowcell with chemistry for 300 cycles and a 150 bp paired end read mode. Sequences were base-called and reads were demultiplexed using bcl2fastq with standard settings.

### Genome assembly

Sequencing data was analysed using the Cambridge University high-performance computer cluster. An assembly-based analysis was carried out on all 73 generated samples using metaGEM^14^, a Snakemakes-based^46^ workflow that allows for the reconstruction of GEMs directly from metagenomic or isolate sequencing samples. In summary, samples were first quality-filtered using fastp^47^ (version 0.20.1) with default parameters. Next, quality-filtered reads were assembled into contigs using MEGAHIT^48^ (version 1.2.9), with the meta-sensitive parameter and minimum contig size of 1 kb. To obtain contig coverage information across samples, each set of quality-filtered short reads was mapped to each generated assembly using bwa^49^ (version 0.7.17). To generate draft bin sets from assemblies, MaxBin2^50^ (version 2.2.7), MetaBAT^51^ (version 2.15), and CONCOCT^52^ (version 1.1.0) were used in combination with the generated contig coverage tables. Following this, the three draft bin sets were consolidated with metaWRAP^53^ (version 1.3.2) to produce a final set of refined and reassembled consensus bins, referred to as metagenome-assembled genomes (MAGs) or single amplified genomes (SAGs) depending on the provenance of the sample. The nine metagenomic samples yielded a total of 32 MAGs. For the 64 isolate sequencing samples, we additionally assembled genomes using shovill (version 1.1.0) (https://github.com/tseemann/shovill), which is an unpublished yet established genome assembly pipeline for isolated genomes. Next, we applied dRep^54^ (version 3.0.0) to dereplicate by choosing the best version of each genome, comparing between metaGEM, shovill, and MEGAHIT outputs, and using dRep’s default genome scoring criteria. Out of a total of 70 generated SAGs, 10 came from MEGAHIT, 28 came from shovill, and 32 from metaGEM. Across the resulting 102 *de novo* sequenced genomes, there was a median completeness of 98.9% (IQR = 17.6%) and a median contamination of 0.82% (IQR = 1.5%) (Supplementary Fig.3b).

### Taxonomy, gene annotations, and identification of mobile genetic elements

GTDB-Tk^55^ (version 2.0.0) with reference data version r207 was employed to assign taxonomic affiliations to the assembled genomes, identifying 4 archaeal and 98 bacterial genomes. Next, the 102 assembled genomes were functionally annotated with the eggNOG^56^ (version 5.0.2) database and eggNOG-mapper^16^ (version 2.1.9). Subsequently, we made use of geNomad^57^ (version 1.5.0) to identify plasmids and bacteriophages across genomes. Furthermore, the ISfinder^58^ database (https://github.com/thanhleviet/ISfinder-sequences) was mapped against the assembled genomes with DIAMOND^23^ (version 2.0.15) to identify ISs with the following parameters: blastp --ultra-sensitive -k 1000. Potentially spurious hits were filtered out using a sequence identity threshold of ≥ 40% for downstream analysis. While this is still not a stringent threshold, it is appropriate given the expected novelty in sequence diversity from such microbes.

### Metabolic modelling

The reconstruction of GEMs was carried out using CarveMe^59^ (version 1.5.1), and gap-filled individually, reflecting the formulation of the cultivation media used. GEMs included an average of 1,513 reactions (SD = 273) and 1,069 metabolites (SD = 162). The auxotrophies function from the ReFramed (version 1.4) metabolic modelling package (https://github.com/cdanielmachado/reframed) was employed to obtain auxotrophy predictions based on flux balance analysis (FBA) simulations for each GEM. Note that while ReFramed is unpublished, it is the underlying library powering tools like CarveMe and SMETANA. These predictions were then compared to the experimentally observed auxotrophic phenotypes. Isolate-based GEMs were further refined by using experimental auxotrophy observations to inform the gap-filling media of each genome. In short, if a given genome was observed to be prototrophic for a given set of amino acids, then these components were removed from the gap-filling medium for that metabolic model reconstruction. Comparing the GEMs auxotrophy predictions against the experimental observations showed that 53 false positive auxotrophy predictions were corrected after the refinement process. Community metabolic interactions were predicted with SMETANA^24^ (version 1.1.0) and carried out by combining individual models from metagenomic and isolate samples that were extracted from the same physical sites. The --detailed flag uses FBA based algorithms^24^ to enumerate all feasible exchanges of metabolic compounds between donors and recipients in a given community, with each exchange having an associated SMETANA score between 0 and 1. Interactions with a score of 0 are not feasible, while interactions with a score of 1 are obligate in order to maintain nonzero biomass flux for the recipient of the metabolic exchange.

### Complementarity of amino acid production capabilities

To determine whether and to which extent auxotrophic bacterial genotypes could complement each other’s metabolic deficiencies, we analysed the amino acid production profiles of 746 auxotrophic isolates that have been sampled from 27 distinct soil microbial communities. The ability of each strain to produce a specific amino acid was represented as a binary vector. This vector consists of 20 binary entries, with each entry corresponding to the presence (1) or absence (0) of the strain’s ability to synthesize a particular amino acid. A value of 1 indicates that the species can produce the amino acid, while a 0 indicates that the species is auxotrophic for that amino acid, meaning it cannot produce it and must obtain it from its environment.

To assess the species’ potential to complement each other’s amino acid production capabilities, we calculated a metric termed ‘complementarity’. For each group of auxotrophic isolates, represented by their respective vectors **v**_**1**_, **v**_**2**_, …, **v**_**n**_ (where n represents the number of auxotrophs in the group), we performed an element-wise comparison using the logical OR operation. This operation was applied across all 20 entries of vectors, resulting in a new vector **r**_**n**_ where each entry is 1 if at least one of the species in the group can produce the corresponding amino acid and 0 only if none of the species can produce it.

A group of strains (**v**_**1**_, **v**_**2**_, …, **v**_**n**_) is considered ‘complementary’ if there is no entry ***j*** such that *r_n,j_* = 0. Mathematically, this means:

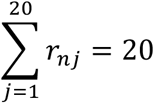

In other words, a group is complementary if each of the 20 amino acids can be produced by at least one of the constituent strains. The total number of complementary groups that meet this criterion of full complementarity (i.e., no 0s in r_n_) is denoted as **N_comp_** The ‘survival complementarity’ for a specific group size n is then calculated as the ratio of **N_comp_** to the total number of possible groups of that size, which is given by the binomial coefficient 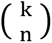, where k is the total number of species under consideration:

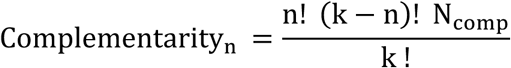

We calculated the complementarity for groups consisting of 2, 3, 4, and 5 auxotrophic isolates. These calculations allowed us to evaluate the degree of complementary in amino acid production within communities of auxotrophs of different sizes.

## Acknowledgments

The authors would like to thank the entire Kost lab (past and present) for useful discussion, as well as Antje Möhlmeyer, Susana Pereira, Björn Reiling, Saskia Schuback, and Eva Limpinsel for technical support. Support by Anja David and Iris Kuhlmann with soil samples collection, Heiko Vogel and Domenica Schnabelrauch (Department of Entomology, MPI-CE) for their help with 16S rRNA sequencing, Sonja Blasche for sharing her DNA isolation protocol and experience, and Bram van Dijk for help with identifying mobile genetic elements is gratefully acknowledged.

This work was funded by the German Research Foundation (DFG: SFB 944, P19, KO 3909/2-1, KO 3909/4-1, KO 3909/6-1, KO 3909/9-1), the Volkswagen Foundation (Az: 9B831), the DAAD German-Egyptian Long-Term program, DAAD End of Scholarship Grants, and Osnabrück University (EvoCell, Female promotion pool).

## Author contributions

Conceived and designed the study: GY and CK; Designed the genome sequencing and modelling: FZ, RG, LO, and KRP; Methodology-Experiments: GY, SD, SG, MI, AS, CK, and SH; Methodology-Sequence analysis: FZ, RG, DW, and VB, Formal Analysis: GY, FZ, LO, and SD; Writing—Original Draft: GY, FZ, KRP, and CK; Writing—Review & Editing: all authors; Visualization: GY, FZ, LO, and CK; Resources, Project Administration, and Funding Acquisition, KRP and CK.

## Data availability

The data supporting this study are available within this article and its Supplementary Information files. The 16Sr RNA gene sequences and the whole genome sequences have been deposited at the European Nucleotide Archive (ENA) (accession code: PRJEB80563). The dataset uploaded to ENA is relevant to Fig. 2c, Fig. 3, and Fig 4d and Supplementary Fig. 9. Assembled genomes and GEMs are available in the GitHub repo linked below.

## Code availability

All code used for genomic analysis, metabolic modelling, machine-learning, statistics, and visualization can be accessed at https://github.com/franciscozorrilla/soil_auxo.

