## Supplemental files for "Obligate cross-feeding of metabolites is common in soil microbial communities"

**Supplementary figures:**

Supplementary figure 1: Locations of soil samples

Supplementary figure 2: Community-level profile of metagenomic samples

Supplementary figure 3: De-novo assembly and binning statistics for 102 genomes

Supplementary figure 4: Phylogenetic tree of 102 de-novo assembled genomes

Supplementary figure 5: Presence and characterization of mobile genetic elements in the 102 isolated and metagenomically assembled genomes

Supplementary figure 6: Size and gene number of the 102 isolated and metagenomically assembled genomes

Supplementary figure 7: Principal component analysis (PCA) of gene copy number based on eggNOG annotations across 102 isolated and metagenomically assembled genomes

Supplementary figure 8: The number of metabolic auxotrophies of a given soil strain correlates negatively with its tendency to grow in experimental cocultures

Supplementary figure 9: Marginally significant correlation between the growth of auxotrophs and the amount of amino acids produced by cocultured prototrophs

Supplementary figure 10: Growth of auxotrophic bacteria depends on an exchange of different essential metabolites with other, co-occurring community members

**Supplementary notes:**

Supplementary note 1: Transcriptional regulators are enriched in genomes with auxotrophic behaviour

Supplementary note 2: The number of plasmid- and virus-encoded genes does not differ between genomes of auxotrophic and prototrophic strains

Supplementary note 3: Depletion of specific genes in genomes with auxotrophic behaviour

Supplementary note 4: Insertion sequences are enriched in genomes of auxotrophic isolates

Supplementary note 5: Amino acid cross-feeding stabilizes the growth of amino acid auxotrophs in a diffusion-based assay of pairwise coculture

**Supplementary methods:**

Supplementary methods 1: Identification of mobile genetic elements

Supplementary methods 2: Amino acid quantification

**Supplementary references**

### Supplementary Figures

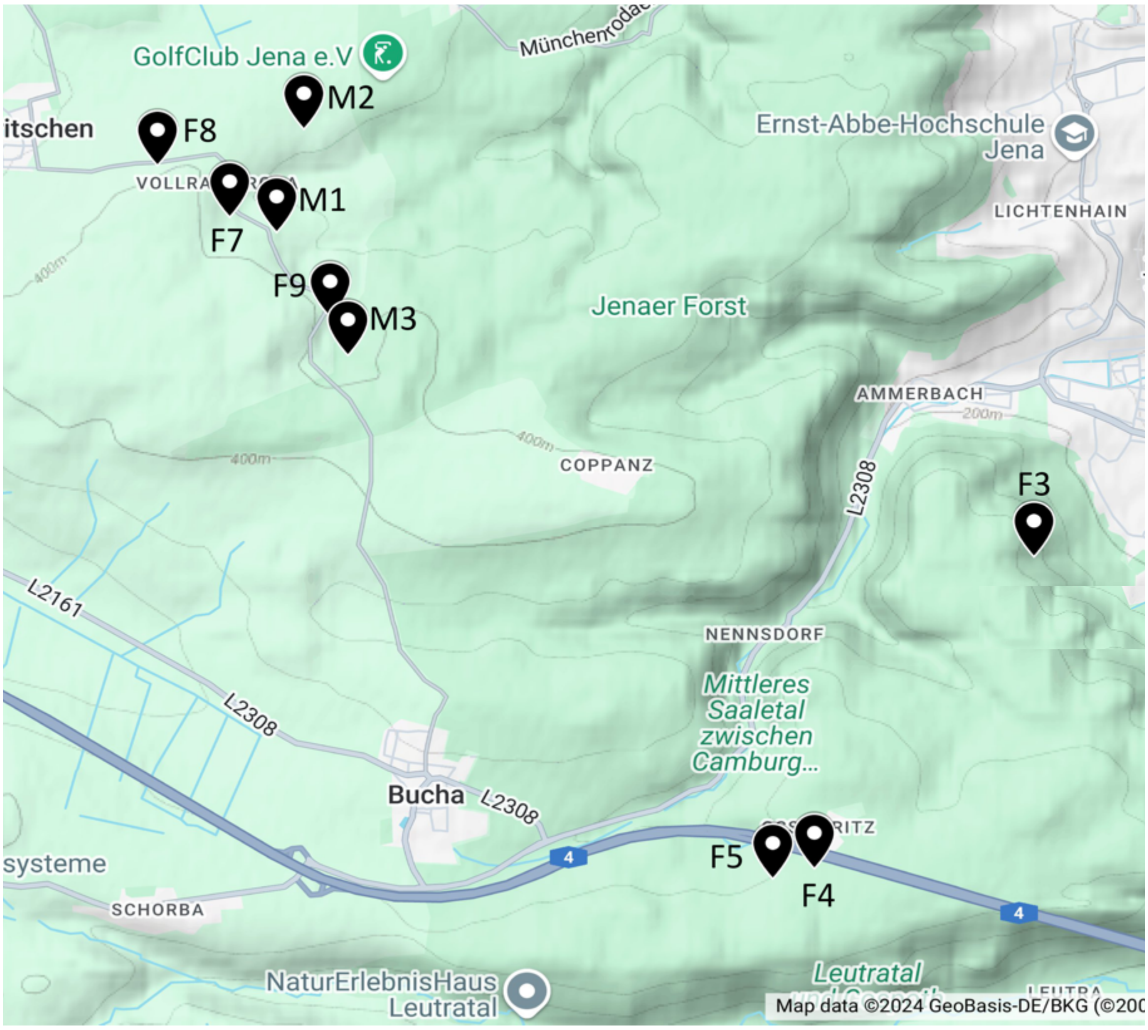


Fig. S1. **Locations of soil samples**. Nine different sites around Jena, Germany, were chosen for collecting soil samples. The coordinates of sites (longitude and latitude) are: M1: 11.499455 50.9139811, M2: 11.5018063 50.9200434, M3: 11.505631 50.9067614, F3: 11.5651667 50.8949266, F4: 11.5461283 50.876613, F5: 11.5425283 50.8760046, F7: 11.4954 50.9148879, F8: 11.48914 50.9179044, F9: 11.5041 50.9090126.

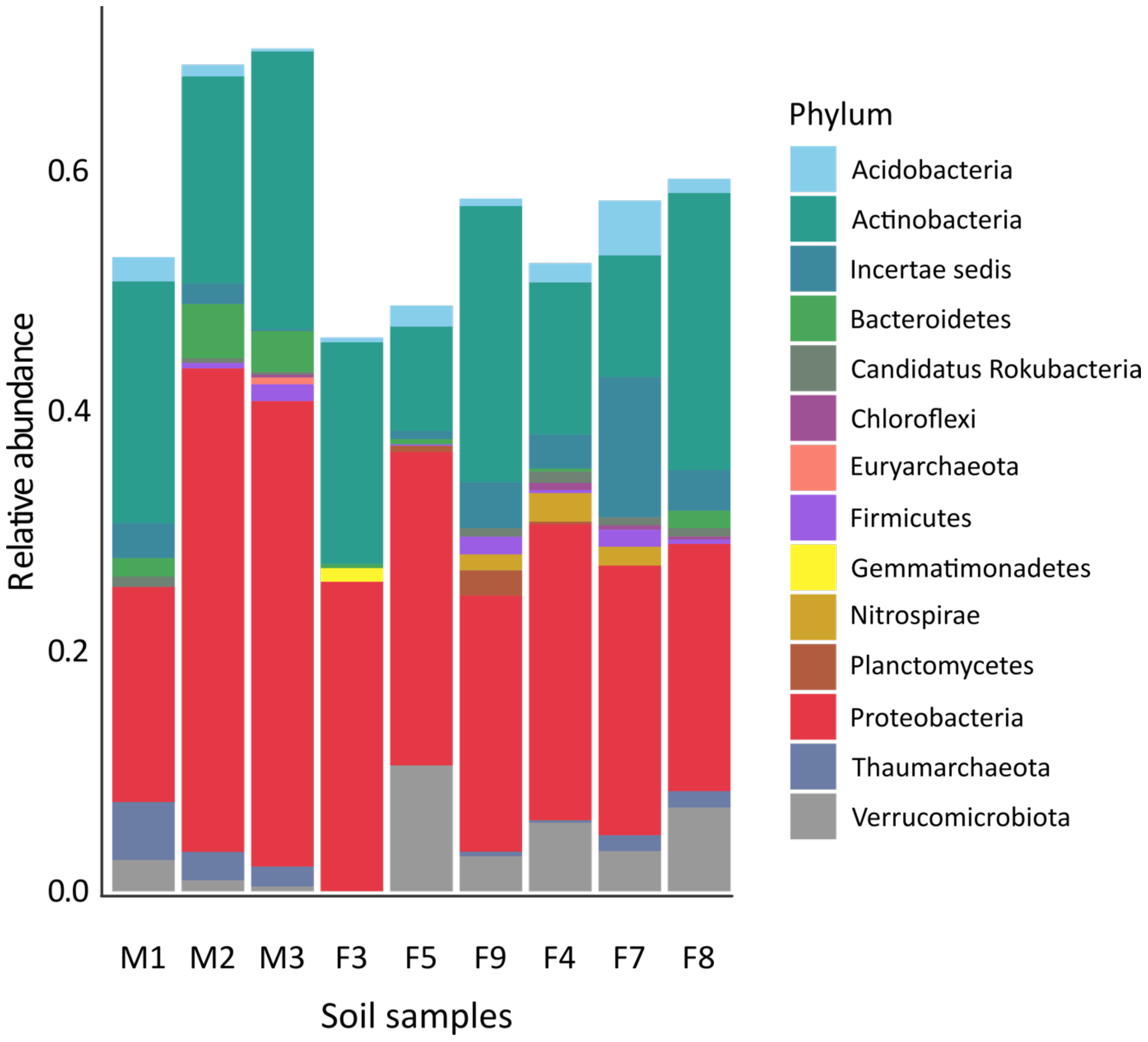


Fig. S2. **Community-level profile of metagenomic samples**. The X-axis shows the nine different soil samples and the Y-axis indicates the relative abundance of different phyla identified.

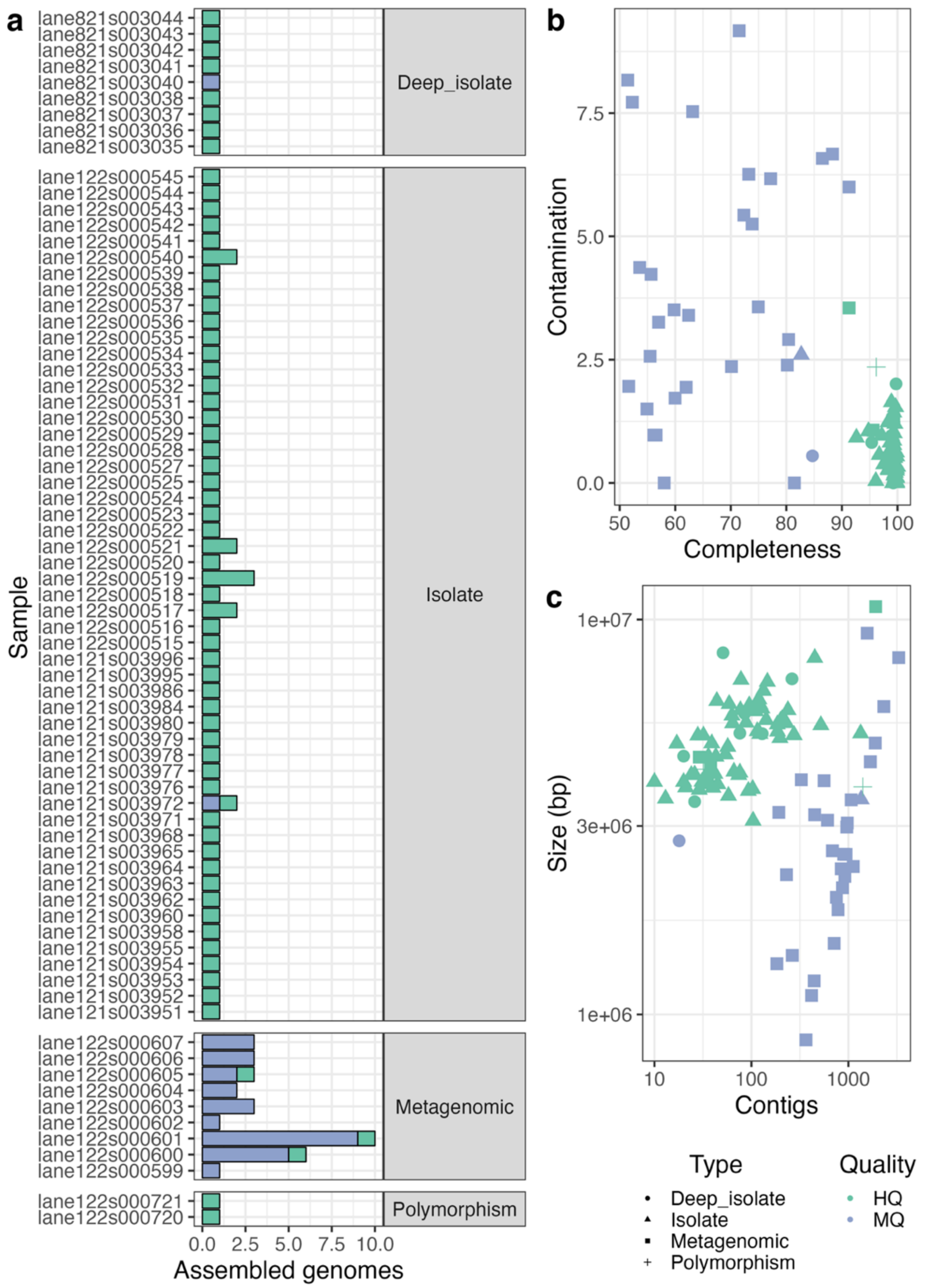


Fig. S3. **De-novo assembly and binning statistics for 102 genomes**. Raw metagenome files were processed using the metaGEM workflow to generate single-amplified genomes and metagenome-assembled genomes. (**a**) Number of genomes reconstructed across sample types. (**b**) Completeness versus contamination estimates. (**c**) Total basepairs and number of contigs across genomes.

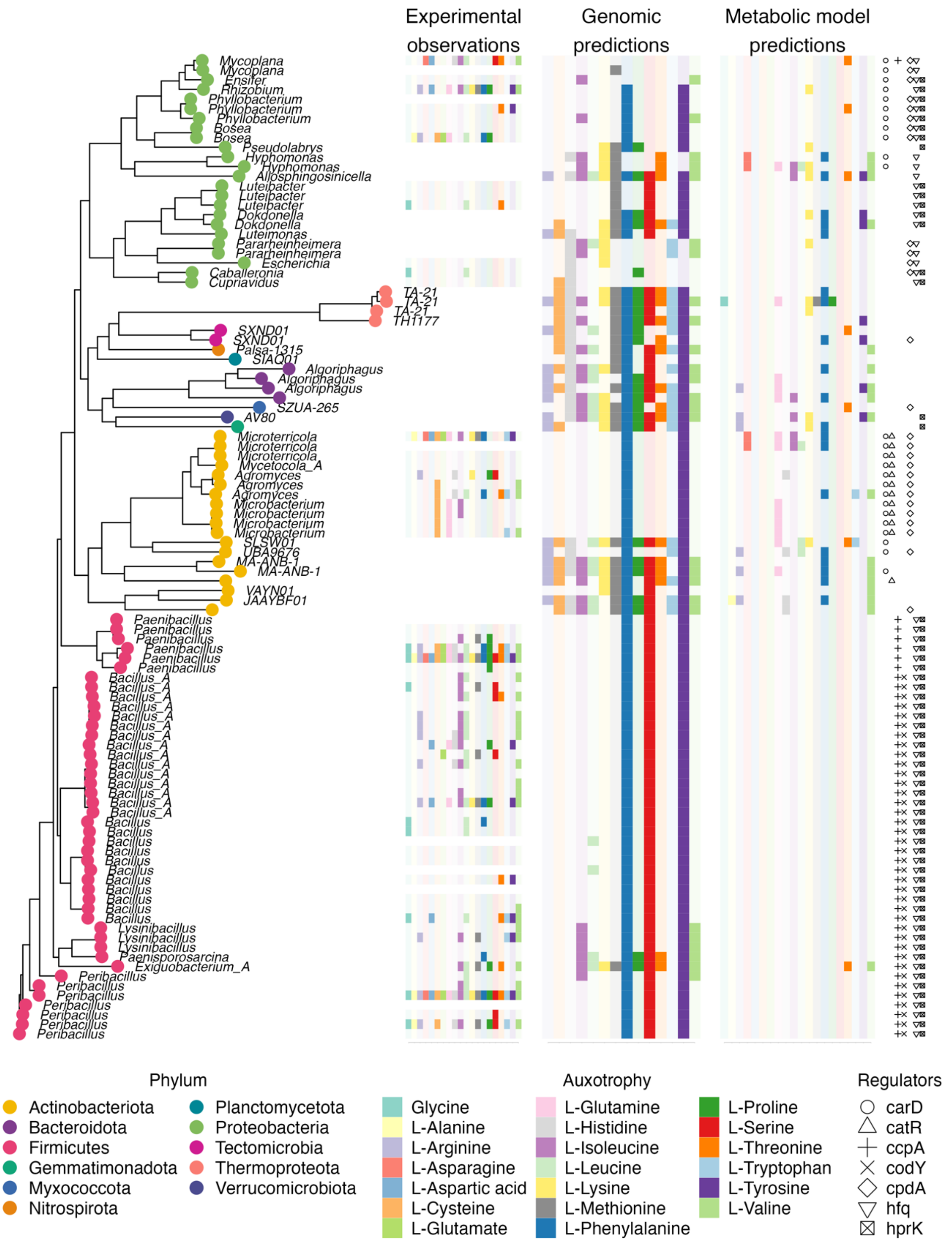


Fig S4. **Phylogenetic tree of 102 de-novo assembled genomes**. Heatmaps show experimental observations, genomic predictions, and metabolic model predictions for amino acid auxotrophies. The legend shows phylum, amino acids, and the presence of metabolic/ transcriptional regulators.

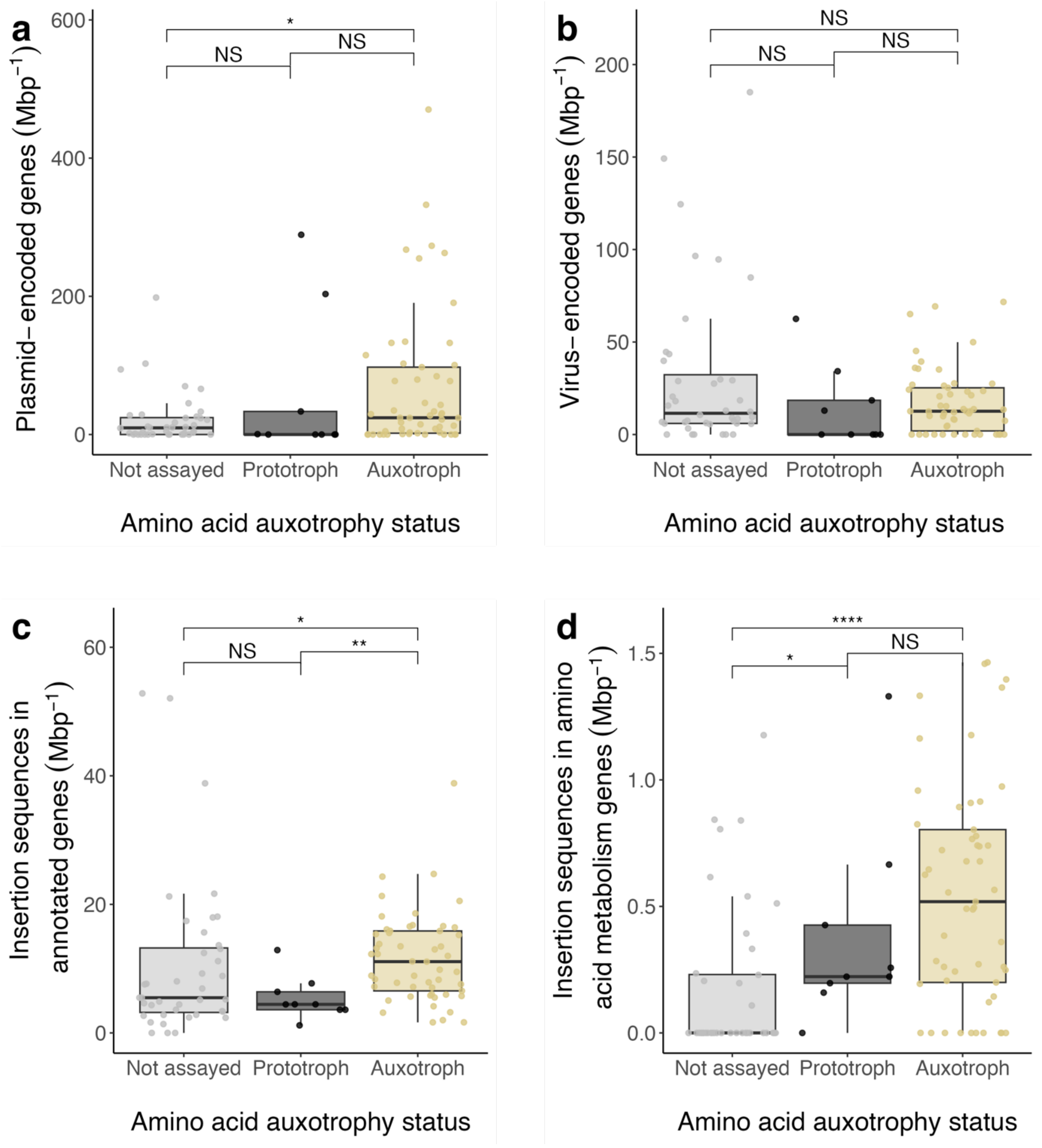


Fig. S5. **Presence and characterization of mobile genetic elements in 102 isolated and metagenomically assembled genomes**. Distribution of (**a**) plasmid and (**b**) virus-encoded genes and insertion sequences across (**c**) all annotated genes and (**d**) amino acid metabolism-related genes per megabasepairs. Genomes of both assembled metagenomes (not assayed) and experimentally verified prototrophic and auxotrophic isolates are included. Boxplots show the median (middle line), lower (25%), and upper (75%) quartiles, and whiskers represent the 1.5 interquartile range. Benjamini-Hochberg corrected *P*-values of Mann-Whitney U tests are shown (NS: not significant, *: $P<0.05$, **: $P<0.005$, ****: $P<0.00005$).

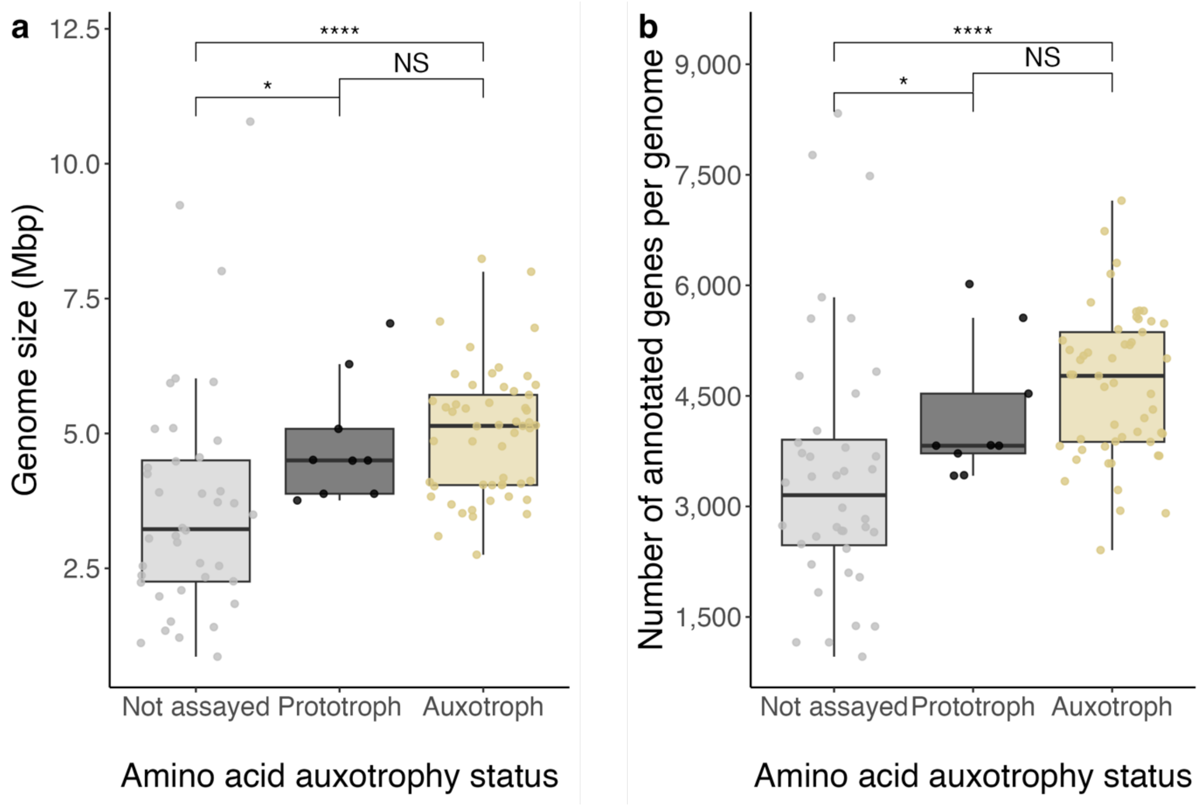


Fig. S6. **Size and gene number of the 102 isolated and metagenomically assembled genomes**. (**a**) Genome size in megabasepairs and (**b**) number of annotated genes per genome across metagenomic (not assayed) and experimentally verified prototrophic and auxotrophic isolates. Boxplots show the median (middle line), lower (25%), and upper (75%) quartiles, and whiskers represent the 1.5 interquartile range. Benjamini-Hochberg corrected P-values of Mann-Whitney U tests are shown (NS: not significant, *: $P<0.05$, **: $P<0.005$, ****: $P<0.00005$).

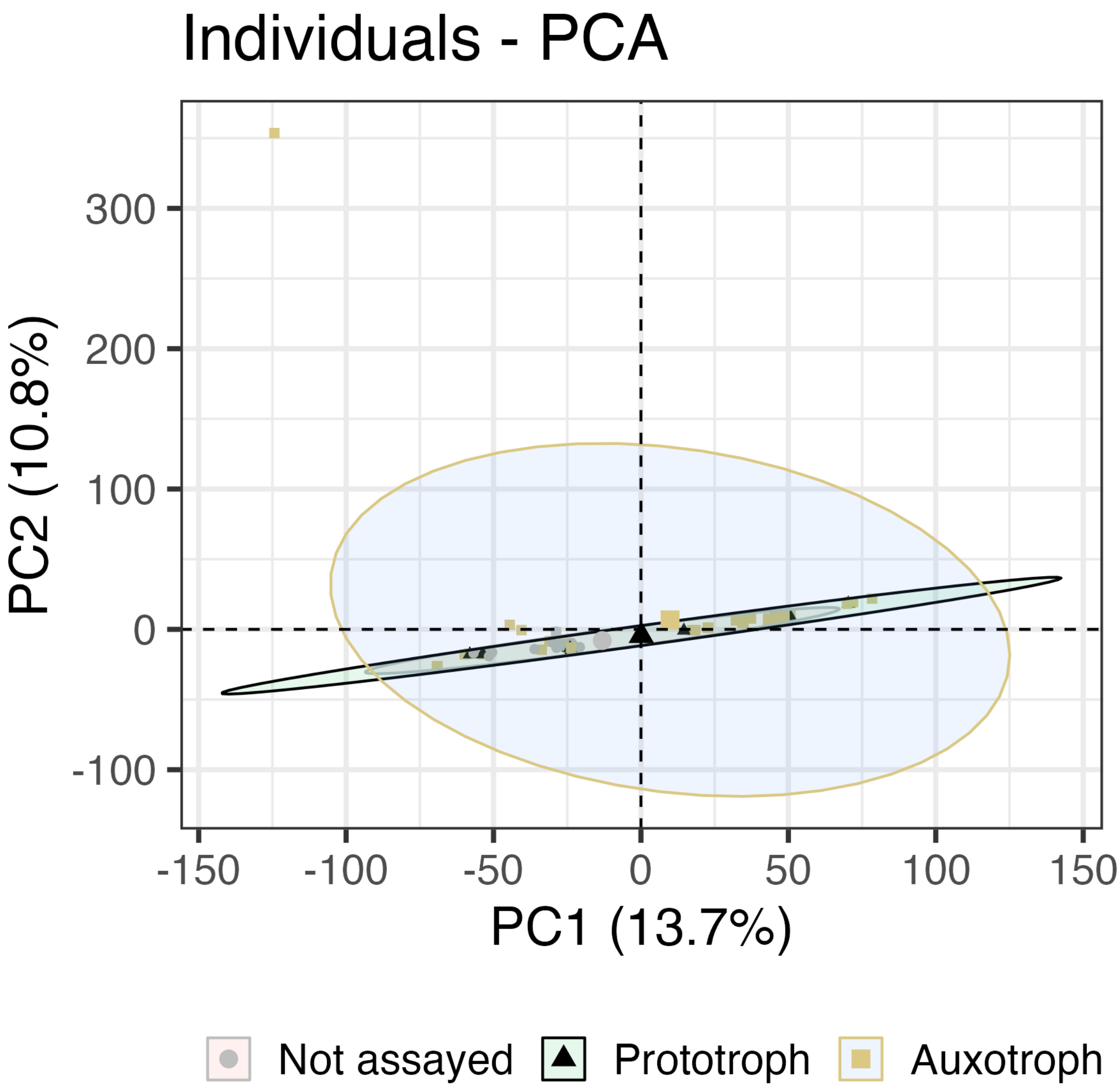


Fig. S7. **Principal component analysis (PCA) of gene copy number based on eggNOG annotations across 102 isolated and metagenomically assembled genomes.**

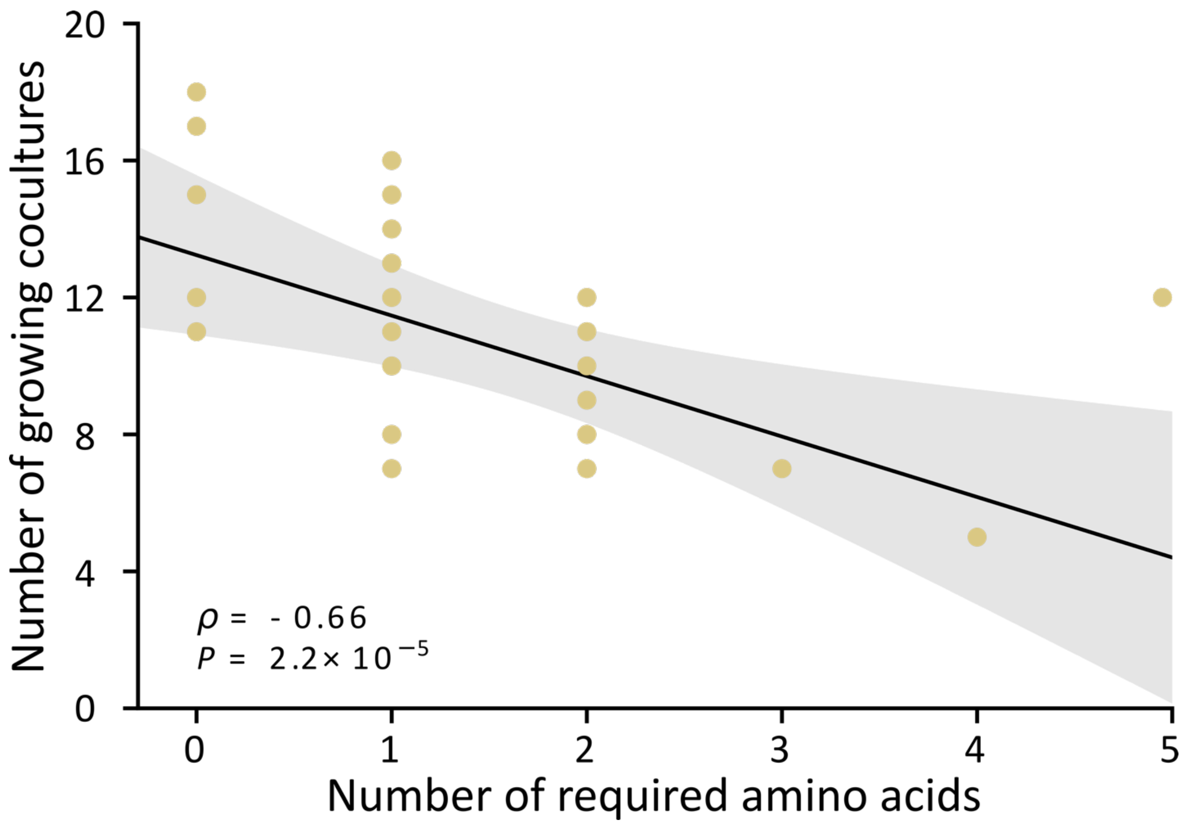


Fig. S8. **The number of metabolic auxotrophies of a given soil strain correlates negatively with its tendency to grow in experimental cocultures**. The number of cases, in which an auxotrophic strain grew in coculture with auxotrophic or prototrophic isolates (y-axis) decreased as the number of amino acids it required for growth increased (x-axis). Results of a Results of a Spearman-rank correlation are shown (n = 23). The black line represents a fitted regression and the shaded area indicates the 95% confidence interval.

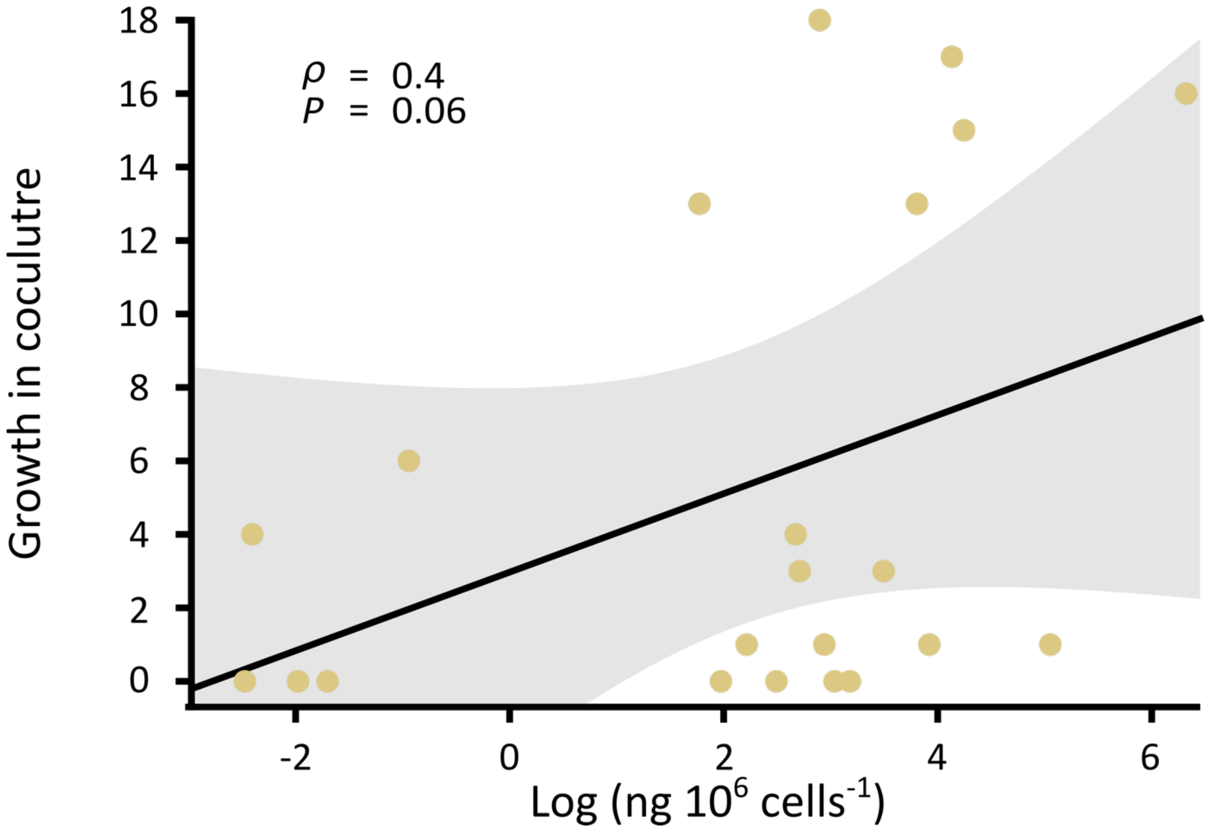


Fig. S9. Relationship between the growth of auxotrophs in coculture with one of 22 prototrophic isolates and the amount of amino acid the prototrophs produced. Coculture growth was calculated as the number of auxotrophs that grew in the presence of the prototroph. Amino acid production rates were determined by growing prototrophic strains in monoculture and analysing the culture supernatant after X h via by LC-MS/MS (for details see Supplementary methods 2). Results of a Results of a Spearman-rank correlation are shown (n = 22). The black line represents a fitted regression and the shaded area indicates the 95% confidence interval.

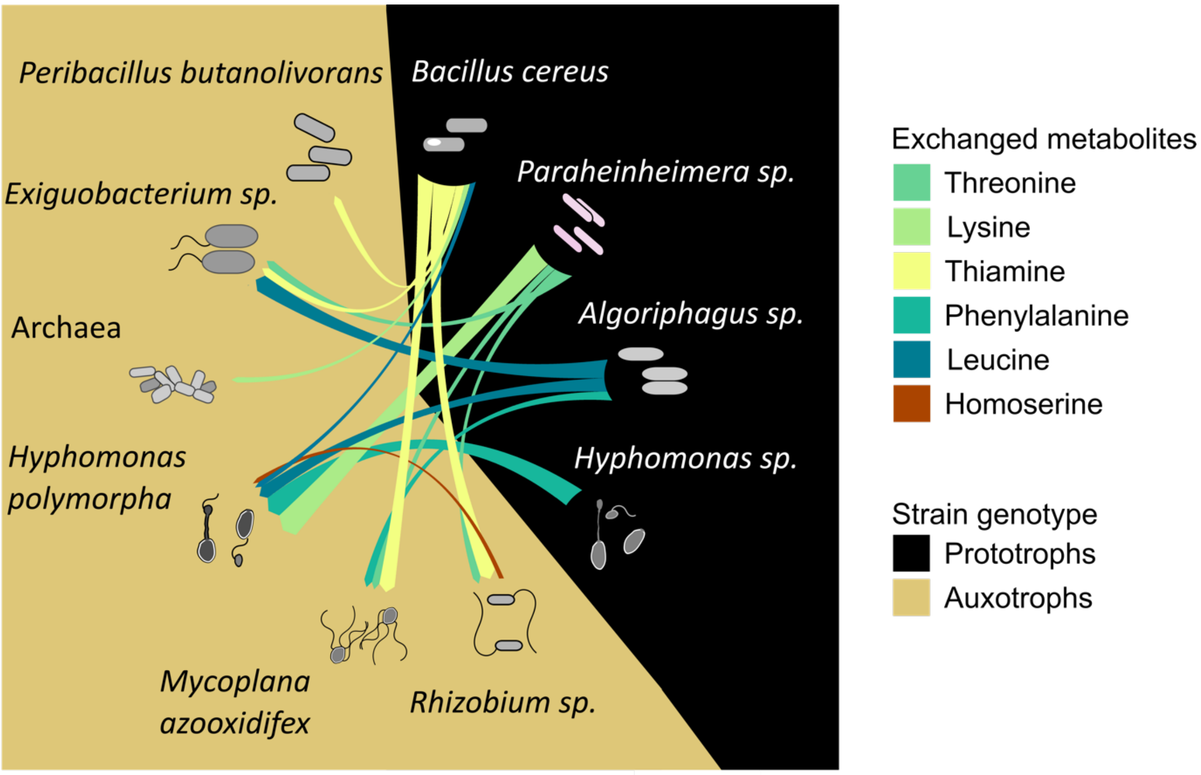


Fig. S10. **Growth of auxotrophic bacteria depends on an exchange of different essential metabolites with other co-occurring community members**. Shown is the exchange of various metabolites (coloured lines) between prototrophic (black background) and auxotrophic genotypes (beige background) as predicted from metabolic modelling of metagenomic samples of a microbial community that has been isolated from site M20. Arrow heads point to metabolite recipients.

### Supplementary Notes

**Supplementary note 1:** Transcriptional regulators are enriched in genomes with auxotrophic behaviour.

When comparing the number and identity of amino acid auxotrophies we observed in our laboratory experiments to the ones that were predicted by the genome analysis, we observed that 33.9% of the leucine, isoleucine, and valine lab-observed auxotrophs carried full biosynthetic pathways. This observation raised the question of whether the evolution of the corresponding auxotrophic phenotypes was driven by transcriptional regulation. Notably, the same pattern has been previously reported from gram-positive bacteria[1], which form the majority in our collection (Fig. 2c). In these cases, the transcriptional regulator CodY[1] could be identified as causal for the observed phenotypic auxotrophies. CodY regulates the operon that is responsible for the production of branched-chain amino acids operon and is highly conserved in Gram-positive bacteria. Interestingly, the presence of exogenous isoleucine in the medium has been reported to impair the growth of gram-positive strains. This is because isoleucine activates the DNA-binding site of CodY [1], thus enabling it to attach to the branched-chain amino acids biosynthetic genes, which in turn causes their repression. The growth of the corresponding strain could be restored by either removing the isoleucine or by an accumulation of mutations that ease CodY's repression[1].

Thus, to test whether CodY can potentially account for some of the observed branched chain amino acids auxotrophies in the gram-positive strains analysed, we verified whether the assembled genomes contained CodY. Interestingly, 90.9% of the analysed cases (40/44 isolated Firmicutes) contained CodY, which may explain the auxotrophic phenotype of these strains. Four isolated Firmicutes, however, lacked CodY (or the gene could not be properly assembled and annotated), which highlights that other factors than transcriptional regulation may drive the evolution of amino acid auxotrophies in these cases.

**Supplementary note 2:** The number of plasmid- and virus-encoded genes does not differ between genomes of auxotrophic and prototrophic strains.

To evaluate whether mobile genetic elements contribute to the emergence of amino acid auxotrophies, we annotated our assembled genomes with the neural network-based plasmid and bacteriophage identification tool geNomad[2] and compared the number of gene hits across genomes. The median count of plasmid- and virus-encoded genes per genome megabasepair was 12.1 (IQR = 46.5) and 12.5 (IQR = 24.0), respectively, with no significant difference between the distributions of genomes with auxotrophic and prototrophic behaviour (Supplementary Fig. 5a and b). However, there was a significant enrichment (Benjamini Hochberg-corrected Wilcoxon rank sum test: $P = 0.045$, $W = 1372$, $n = 93$) in the count of plasmid-encoded genes per megabasepair for auxotrophic genomes (median = 24.2, IQR = 95.6, n = 53) compared to metagenome-assembled-genomes (median = 9.62, IQR = 24.5, n = 40) (Supplementary Fig. 5a).

Furthermore, plasmid-encoded genes were generally enriched in cultivated isolates compared to MAGs (Wilcoxon rank sum test: $P = 0.035$, $W = 1410$, $n = 102$); although this is likely related to methodological limitations in recovering plasmids from metagenomic samples. There was also evidence of amino acid metabolism genes being present on plasmids. Intriguingly, one *Rhizobium sp900472625* (lane122s000521_bin.2.p) strain provided strong evidence that up to 49.9% of all its amino acid metabolism genes were localised on a plasmid. Furthermore, this microbe could not be separated from its putative metabolic partner *Bacillus wiedmannii* (lane122s000521_bin.1.s), suggesting the plasmid might be functionally involved in mediating the obligate metabolic relationship between both strains. In these cases, loss of the corresponding plasmid can be a rapid way to generate auxotrophic genotypes in a subpopulation of cells[3]. If the corresponding profile of metabolic auxotrophies is beneficial under the current environmental conditions, the auxotrophic variants would increase in frequency relative to the plasmid-containing prototrophic strain. If, however, conditions switch back to the previous state, the relative frequencies of prototrophic, plasmid-bearing cells is likely re-adjusted by natural selection. Finally, there was scant evidence of virus-encoded genes that are related to amino acid metabolism (median = 0, IQR = 0): only 21 such genes were found across 12 MAGs. Notably, one unknown metagenomic species (genus UBA9676 within the family Streptosporangiaceae) stood out for having 1,377 viral genes across ~780 kbp, representing 8.5% of its total genome content. Overall, these findings suggest that viruses are likely less relevant than plasmids for encoding amino acid biosynthesis and potentially causing auxotrophic behaviour when lost.

**Supplementary note 3:** Depletion of specific genes in genomes with auxotrophic behaviour.

Gene annotations were used to identify variation in gene copy numbers across genomes. In this way, genes could be identified that were particularly enriched or depleted genes in auxotrophic and prototrophic genomes. Genes that were globally enriched included genes involved in replication and repair (*yaoA, dinG, mutL, mutS*), protein degradation and translocation (*hslU, hslV, yajC*), and regulation of metabolism (*hprK,hfq*). Notably, depleted genes were involved in amino acid metabolism and transport (*glnA2, glnA4, apeB, pipch2, gadB, aspC, ilvJ, proX*), peptidases (*pepO, rip1, hofD*), or transcriptional or metabolic regulators (*abaB2, cpdA, catR, carD*). However, a principal component analysis (PCA) of gene copy number across genomes did not show good separation between genomes with auxotrophic and prototrophic behaviour (Supplementary Fig. 7).

**Supplementary note 4:** Insertion sequences are enriched in genomes of auxotrophic isolates.

To further investigate the functional role insertion sequences (ISs) play in the evolution of metabolic auxotrophies, we inspected the intersection of IS hits and eggNOG gene annotations, yielding a total of 5,549 annotated genes with evidence of ISs. Out of these, 48.9% were related to replication and repair (L), 12.1% had unknown function (S), 9.1% were related to transcription, 5.7% had no COG category assigned (-), 3% were related to inorganic ion transport and metabolism (P), and 2.2% related to amino acid transport and metabolism (E). The number of annotated genes with evidence of ISs per megabasepair was significantly greater in genomes of auxotrophic strains (median = 11.1, IQR = 9.32, n = 53) compared to genomes of prototrophic strains (median = 4.44, IQR = 2.79, n = 9) (Benjamini Hochberg-corrected Wilcoxon rank sum test: $P = 0.009$, W = 90, n = 62) (Supplementary Fig. 5c).

Strikingly, IS hits were enriched in Firmicutes (median = 238, IQR = 297, n = 44) relative to Actinobacteria (median = 16.4, IQR = 82.2, n = 19) (Benjamini Hochberg-corrected Wilcoxon rank sum test: P = 0.0002, W = 130, n = 63). This observation corroborates previous studies that reported certain phyla may act as IS reservoirs, thus potentially playing an important role in transcriptional regulation and transport[4]. No significant differences was detectable when the number of ISs per megabasepair was compared between isolates and metagenome-assembled genomes (Wilcoxon rank sum test: P = 0.418, W = 1198, n = 100).

**Supplementary note 5:** Amino acid cross-feeding stabilizes the growth of amino acid auxotrophs in a diffusion-based assay of pairwise coculture.

We asked whether the propensity of auxotrophic recipients to grow depended on the amount of amino acids that prototrophic donors provided in coculture. To address this, we quantified the total concentration of amino acids that was detectable in the supernatant of monoculture of each prototroph that have been used in the coculture experiment (Supplementary methods 2). The results of this experiment provided weak statistical support for the hypothesis that the amino acids that are externalized by prototrophs enhance the growth of the cocultured auxotrophs (Pearson correlation: $r = 0.4$, $p = 0.06$, $n = 22$, Supplementary Fig. 9). A possible explanation for this is that many of the tested auxotrophs required multiple different amino acids to grow. Consequently, each of these strains required a very specific mixture of metabolites, which likely blurred the statistical relationship between both parameters.

### Supplementary methods

**Supplementary methods 1.** Identification of mobile genetic elements.

We used geNomad[2] (version 1.5.0) with default parameters to identify plasmids and bacteriophages across genomes. The number of plasmid and virus-encoded genes was counted across assembled genomes with Rstudio (version 4.1.0) and tidyverse (version 2.0.0).

**Supplementary methods 2.** Amino acid quantification.

Precultures of 18 prototrophic genotypes that were previously used in the cross-feeding experiment (Fig. 4a) were inoculated into MMAB without supplements. After three days, cells were collected during their early stationary phase by centrifugation and their supernatant was filtered using a pore-size of 0.22 µm (PALL Corporation, Multi-Well Teller). Cell-free supernatants were stored at -20 °C until further use. To normalize the data, the number of colony-forming units (CFUs) of each strain was determined by serially diluting 20 µl of the culture, drop plating 15 µl of each dilution on a solid MMAB medium, and incubating the plates for three days at 30 °C.

Mixtures of all 20 amino acids were prepared at different concentrations to generate a calibration curve. Both the previously collected cell-free supernatants and the amino acid mixtures for the calibration curve were derivatized as described elsewhere[5]. In detail, 50 µl of each supernatant and the calibration mixtures were individually added into PCR tubes (Starlab) that each contained 10 ml of sterilized deionized water, 90 µl MMAB broth, and 1 ml of 1M NaOH. To this mixture, a 2.5 µl aliquot of norleucine (Alfa Aesar) was added as internal standard. Derivatization was performed by adding 50 µl of dansyl-chloride solution (Merck), which was freshly prepared by adding 10 mg of dansyl-chloride to 1 ml of acetonitrile (AppliChem). Next, tubes were incubated at 80 °C for 40 minutes in a PCR machine (Biometra - TAdvanced). After incubation, 5 µl of 351 mM formic acid solution was added to each tube and the samples were stored overnight at 4 °C. The next day, samples were centrifuged for 20 minutes at 4,200 rpm and 100 µl of the supernatant were transferred into a sterile 96-well deep-well plate (Greiner), covered with MS-Sealing foil (Clearline Sealplate – Kisker Biotech G040), and subjected to mass spectrometry (Sciex, ABSciex Qtrap 5500). Chromatography was carried out using a Shimadzu HPLC system and the separation of the phases was performed with an Accucore RP-MS 150 3 2.1, 2.6 mm column (Thermo Scientific). The mobile phase used was formic acid 0.1 % in 100 % water and 80 % acetonitrile, its flow rate was 0.4 ml $min^{-1}$, and the injection volume was 1 ml. The liquid chromatography was coupled to a ABSciex Q-trap 5500. The curtain gas had a pressure of 40 psi, collision gas was set to high, ion spray voltage was set to 2.5 keV, the temperature was 550 °C, ion source gas (1) was set to 60 psi, and the ion source gas (2) was set to 70 psi [5].

The data obtained was analysed using the SciexOS software (Sciex, version 3.1). Norleucine was used as an internal standard in each sample to compensate for variations in dilution, evaporation and/or decomposition. A calibration curve was used to determine the linear detection range and the "response factor" of the mass spectrometer.
